# Comprehensive Analysis of Molecular Characteristics, Clinical Signifificance, and Cancer Immune Interactions of Patients by Anoikis-Related Genes in LUAD Combined with Single-cell Data

**DOI:** 10.1101/2023.05.15.540748

**Authors:** Weijie Yu, Zhoulin Miao, Julaiti Ainiwaer, Bingzhang Qiao, Kawuli Jumai, Ilyar Sheyhidin

**Author notes:** Correspondence should be addressed to Ilyar Sheyhidin.

## Abstract

**Background:** Lung adenocarcinoma(LUAD) is the most prevalent subtype of lung cancer today. There is a close relationship between Anoikis related genes(ARGs) and tumor prognosis, drug susceptibility, and tumor microenvironment(TME).

**Method:** We calculated differential expression genes using downloaded Anoikis genes and selected genes of prognostic value. Consensus clustering analysis was used and characterized between different clusters. Differences between the different groups were also explored. Risk scores and Nomogram with predictive prognostic functions were established. Immune status and drug sensitivity were also assessed between different risk groups. Single-cell data were downloaded to compare the expression profiles of selected genes, and immunohistochemical results of selected genes were also downloaded to corroborate the reliability of the manuscript.

**Result:** Two clusters were identified on the basis of related gene expression. We analyzed the survival time, functional enrichment between the two groups and found significant differences between the two clusters. Significant relationships were found between the different clusters and clinical variables. group B had a significantly lower KM curve than group A, as well as a significant enrichment in multiple tumor functions. A risk score with prognostic value was established. The risk score was found to have a high predictive value for prognosis and was an independent prognostic factor. Combined with clinical variables, a Nomogram was established and found to be an accurate predictor of patient prognosis. There were significant differences in immune status between the different risk groups. Patients in the low-risk group were significantly better treated than those in the high-risk group. Finally single cell data confirmed the expression of the selected genes. Also, the immunohistochemical results helped us to confirm the selected genes have increased expression in tumor tissue.

**Conclusion:** In conclusion, this paper reveals the role of ARGs and immune status, drug susceptibility, and prediction of prognosis in LUAD. Also, an accurate prognostic prediction model was established based on genetic.

## INTRODUCTION

Lung cancer is recognized as the number one malignancy in the world today. According to statistics, the annual number of new cases accounts for 11.4% of all cancer records, second only to breast cancer at 11.7%, and the mortality rate accounts for nearly 20%, which is also the highest mortality rate of neoplastic diseases [1]. Among them, lung adenocarcinoma(LUAD) is the most prevalent subtype of lung cancer(LC). Because symptoms of early stage LC are not apparent, it is not easy to attract patients’ attention, so the best time for surgical treatment is often missed once it is diagnosed. For patients with advanced stages, chemotherapy, targeted therapy, and immunotherapy are indispensable treatments. Due to the alteration of tumor microenvironment, it not only affects the drug efficacy, in addition, but also leads to immune escape. Under the current problem, exploring other mechanisms of induced cell death becomes the essential to solve the problem.

Impaired apoptosis is a characteristic of tumor cells. The imbalance of apoptosis empowers cells to resist drug-induced apoptosis.Among the many forms of apoptosis, Anoikis is considered as a novel mode of apoptosis that can perform protective and stabilizing tissue functions [2]. When cells are separated from the extracellular matrix(ECM) during transfer, this triggers apoptosis initiation and prevents abnormal cellular proliferation, thus achieving protection [3, 4]. However, tumors do not need to be attached to the ECM in order to survive [5].This non-anchored growth grants significant independence to the tumor cells and generates resistance to apoptotic genes through the associated mediators. As a result of this acquired resistance to apoptosis, it is referred to as Anoikis resistance. The result is that the adherent cells continue to survive or proliferate with other ECM, or metastasize to distant sites[6]. According to the results of existing studies, genes involved in Anoikis have a close relationship with tumor development. In patients with LC, FAIM2 and PDK4 are significantly correlated with poor prognosis and poorer chemotherapeutic drug effects [7, 8]. However, the present studies have only focused on individual genes and lack a more comprehensive and more systematic evaluation. Chemotherapy and immunotherapy are indispensable therapeutic options for patients with advanced stages. However, treatment effectiveness and tumor microenvironment(TME) are clearly correlated [9]. In pathological conditions, tumor cells reverse the tumor microenvironment and promote the suppressive effect of related immune cells resulting in the loss of surveillance and clearance of tumor cells [10]. As a novel mode of apoptosis, the relationship between Anoikis genes and drug sensitivity and TME is unclear. Therefore, in this study, we comprehensively assess the impact of Anoikis genes on drug sensitivity and TME in patients with lung adenocarcinoma. We will also develop a predictive model in combination with clinical features to assist in the assessment of patient prognosis.

## MATERIALS AND METHOD

### Data collection

We obtained the anoikis genes by two websites(https://www.genecards.org and https://maayanlab.cloud/Harmonizome). Setting the relevance score > 0.4, we obtained a total of 640 genes. We obtained the LUAD patient cohort and corresponding clinical information through the TCGA database (https://portal.gdc.cancer.gov/), copy number variation. GSE26939 data were obtained through the GEO website (www.ncbi.nlm.nih.gov/geo/). Patient data with lack of survival data or significant clinical information were further excluded. To obtain differentially expressed genes (DEGs) of anoikis genes in LAUD, we took the intersection of anoikis genes and TCGA-LUAD data and performed differential analysis of intersecting genes between normal and tumor tissues using the limma package in R package (|log2-fold change (FC)| ≥ 1, p-value <0.05). The TCGA-LUAD and GEO data were normalized and combined to extract gene expression. We performed unicox regression analysis of DEGs, and genes with prognostic significance were selected and analyzed for follow-up (p<0.05).

### Consensus Clustering Analysis of ARGs

Consensus clustering using the k-means algorithm is used to define different patterns of anoikis. The number and consistency of the clusters were established by the consensus clustering algorithm available in the “ConsensuClusterPlus” package [11]. 1000 iterations were performed to ensure the stability of these classes. A CDF plot was used to help determine the optimal K values. PCA analysis was also used to directly evaluate the K values. The GSVA, GSEAbase package was then used to complete the single-sample gene set enrichment analysis (ssGSEA) analysis to calculate the differences in immune cell subtypes between groups.

### Functional enrichment analysis between ARG cluster groups

Gene set enrichment analysis (GSEA) was then performed to identify the most signifificantly enriched pathways between the two clusters through R software, “org.Hs.eg.db,” “clusterProfifiler,” and “enrichplot” packages. Kyoto Encyclopedia of Genes and Genomes (KEGG) and Gene Ontology (GO) analyses were exploited to investigate the most signifificantly enriched pathways and biological processes of the DEGs using R software, “clusterProfifiler”, “GSVA”,“GSEABase” package.

### Construction of a risk model related to anoikis

we performed univariate COX regression analysis and least absolute shrinkage and selection operator (LASSO) regression to gain the anoikis related genes for predicting survival and prognosis of GBM. The calculation formula of risk score is shown below: Risk score = ΣCoef (ARGs) * Exp (ARGs), where Exp (ARGs) is the relative expression of the candidate ARGs, and Coef (ARGs) is the regression coefficient. Kaplan-Meier analysis was used to compare the OS between the two groups with the “survival” and “survminer” R packages. ROC curves were used to assess the accuracy of the model. We further aggregated the model and clinical characteristics (age, sex, stage) and performed multi cox regression analysis to determine whether the model was an independent prognostic factor. We analyzed the expression patterns of selected genes at the protein level through the Human Protein Atlas (HPA) website (https://www.proteinatlas.org/). Finally, data were obtained through the single cell database TISCH2 (http://tisch.comp-genomics.org/).

### Construction of Nomogram Based on anoikis and Clinical Characteristics

By “rms”package, risk scores and other clinical characteristics of patients(Age, Gender, Stage) are aggregated to develop Nomogram in order to provide valuable clinical predictions for LAUD patients with, particularly, 1-, 3-, and 5-year OS[12]. Next, calibration curves and decision curves analysis (DCA) were plotted to verify the clinical validity of the established Nomogram.

### Analysis of immune characteristics between risk score group

The CIBERSORT algorithm was used to assess the level of immune cell infiltration between the different risk groups. We then used the selected genes to assess the relationship between and immune cell subtype. In addition to this, the relationship between immune cells was also evaluated. We used the ESTIMATE algorithm to assess TME score in LUAD patients.

### Correlation of Drug sensitivity and risk score

Drug response information was collected in the Genomics of Drug Sensitivity in Cancer (GDSC), and drug response was compared between high and low risk groups using “oncopredict” package, and box plots were drawn for the meaningful comparison groups[13]. Due to the close relationship between immune checkpoint expression and effect of immune checkpoint inhibitors (ICIS), we compared the expression of 47 immune checkpoints[14,15]. Tumor immune dysfunction and exclusion (TIDE) scores (http://tide.dfci.harvard.edu/) were calculated for each LUAD sample in the cohort, and the difference in TIDE scores between high first risk groups was determined_[16]_.

### Statistical Method

All statistical analyses were performed using R software (4.2.1). Kaplan-Meier (K-M) analysis was applied to assess survival difffferences between the two groups. Reliability of the ARGs models was tested using receiver operating characteristic (ROC) analyses. P-value< 0.05 was considered statistically signifificant (*p-value< 0.05; **p-value< 0.01; ***pvalue< 0.001).

## RESULTS

### Genetic alterations and expression of ARGs in patients with LUAD

In this article, a total of 640 ARGs were included(Table S1). First, we compared the expression of ARGs in LUAD samples and normal tissues. A total of 129 genes were significantly different between normal and tumor tissues (Figure 1A, Figure 1B, Table S3). Next, we explored the effect of ARGs on patient survival (OS). We did a uniCox and construct Forest plot of the differential genes for ARGs and showed that a total of 26 genes showed significant difference (Figure 1C, Table S4). We visualized the relationship between ARGs and prognosis of LUAD patients using network plots (Figure 1D, Table S5).

**Figure 1.**
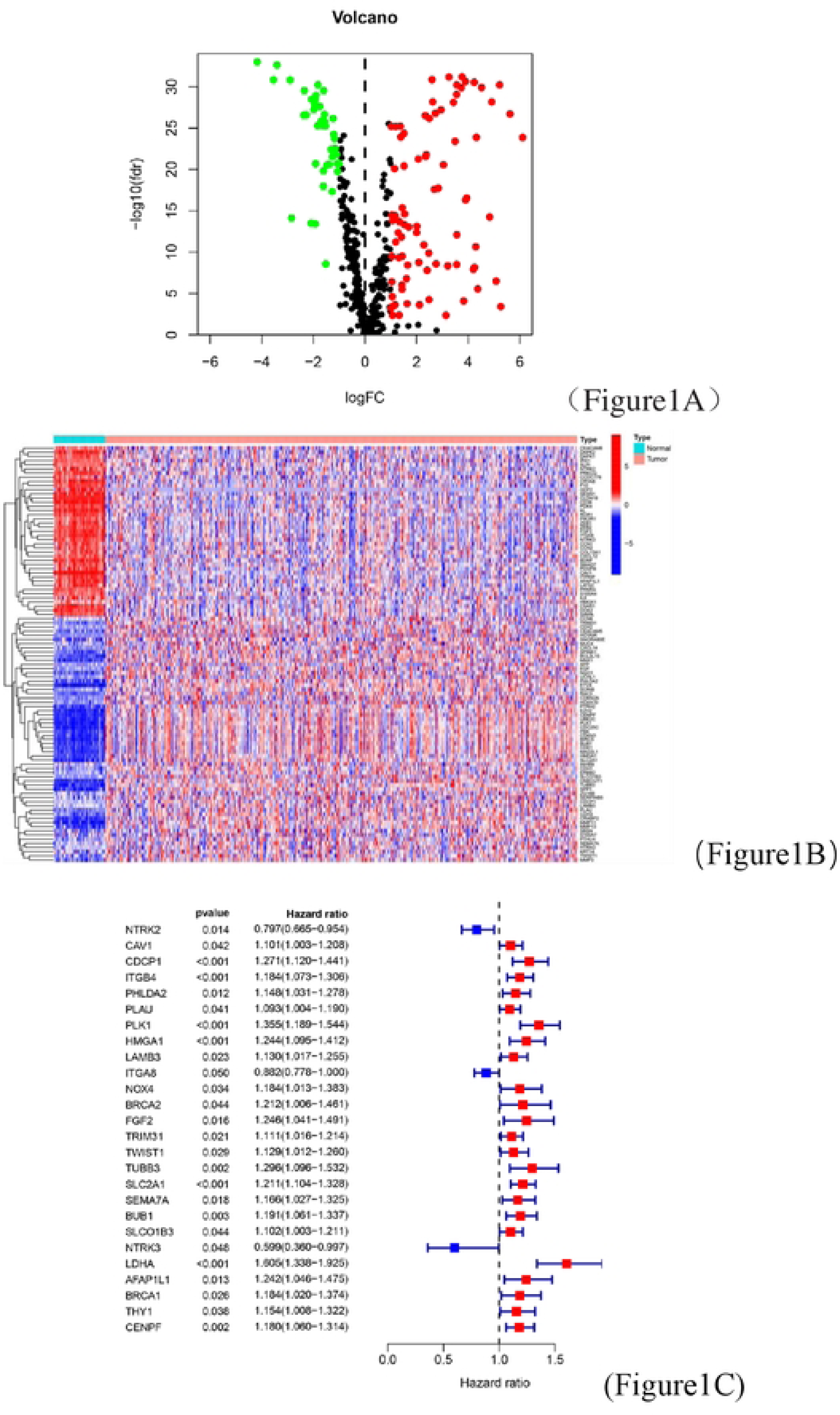
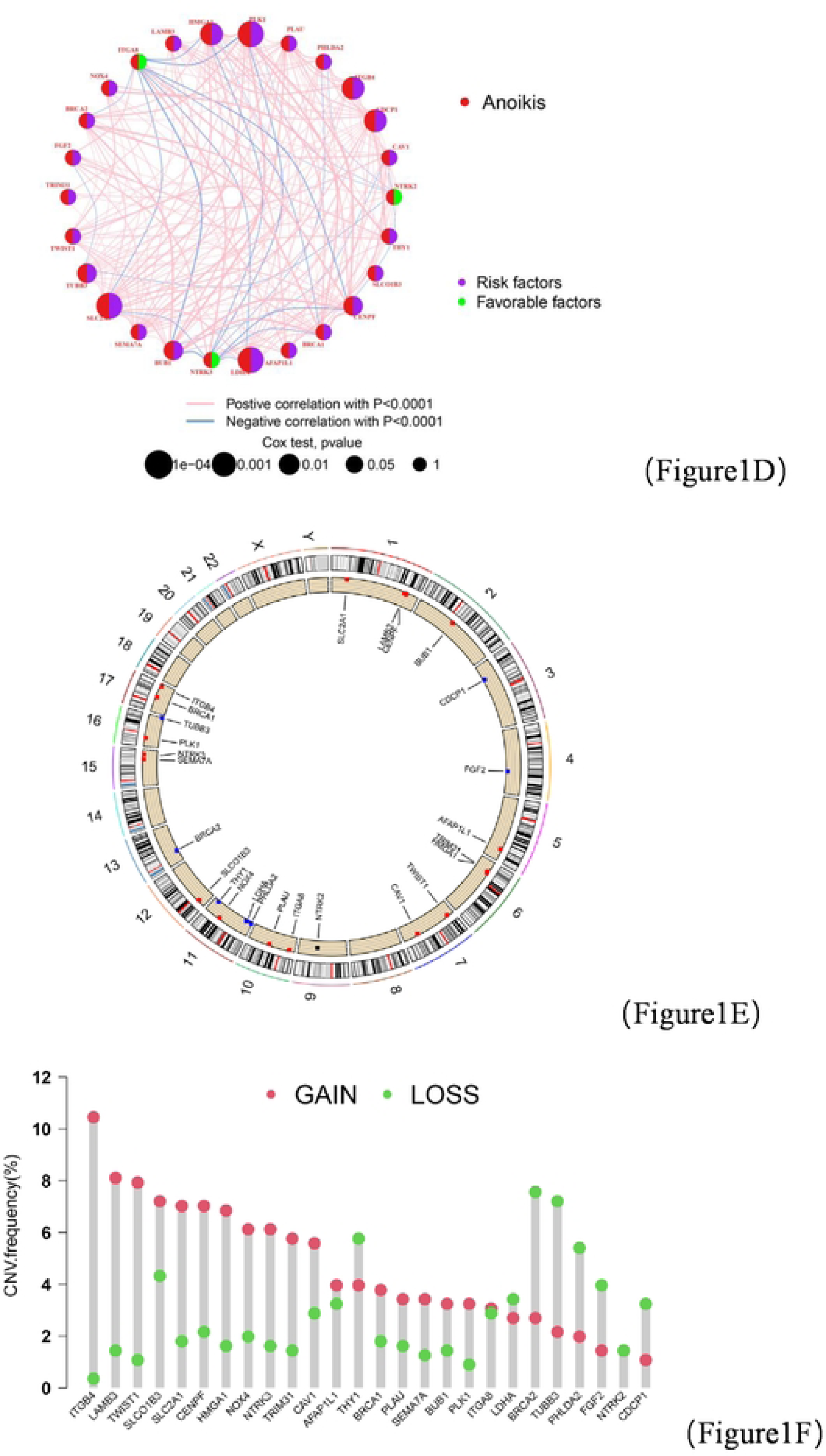
Identification of genetic alterations and loss of apoptosis-related gene expression in LUAD patients. (A) Volcano map of differential genes. (B) Heat map of differential gene expression in normal and tumor tissues. (C) univariate Cox regression analysis of differential genes. (D) Prognostic network relationship map. (E) Circus plots of chromosome distributions of AAGs. (F) Frequency of CNV gain, loss and non-CNV in loss of apoptosis genes (p < 0.05 *; p < 0.01 **; p < 0.001 ***)

The size of the circles in the figure represents the P-value of univariate analysis of genetic prognosis. Blue represents risk factors and green represents favorable factors. The colors of the connecting lines between the dots represent positive and negative effects, respectively. Figure 1E demonstrates the mutation sites and alterations of genes on chromosomes. In addition, we explored the incidence of CNV mutations, which showed that 26 genes showed significant CNV alterations (Figure 1F, Table S19). The above results indicate that there are significant differences in the genomic background and expression levels of ARGs between tumor tissues and normal tissues, suggesting that it is the ARGs that may play an important role in LUAD.

### Differences in subtypes, clinical variables, and TME of Anoikis

Complete information of these patients was listed in Table S2. We used consensus clustering analysis to classify genes with prognostic significance. Meanwhile, the optimal K value is selected by referring to the CDF curve. So we divided the samples into 2 clusters(Table S6). After grouping the patients, we compared the overall survival (OS) of the patients, plotted K-M curves and compared the survival difference between the two groups (p<0.001) (Figure 2C). PCA analysis also showed significant differences between the two groups (Figure 2D). As in Figure 2E, we compared the gene expression differences between the two cluster of patients and found that most of the genes were differentially and highly expressed in ARG cluster B. We plotted heat maps to reveal the relationship between gene expression and clinical characteristics. In the results we found a significant relationship between Anoikis genes and clinical variables in patients (Figure 2F). As showed in Figure 2G, most of immune sub-cell were different between two clusters(Table S7).Finally, we performed an enrichment analysis of these genes and found significant differences between the two clusters. Cluster B was mainly enriched in the biological processes of tumor-related diseases (small cell lung cancer, prostate cancer) (Figure 2H, Table S9). GO analysis was performed between the clusters, and we could clearly find significant differences between cluserB and clusterA (Figure 2I,Table S8). clusterA was hardly significantly enriched in BP, CC, and MF. Then we performed GSEA analysis on the two clusters. By comparing Figure J-K, we found that clusterA was active in arachidonic acid metabolism. In contrast, cluster B was active in other pathways(Table S10). Comparing Figure L-M, we found that clusterA was hardly expressed active(Table S11).

**Figure 2.**
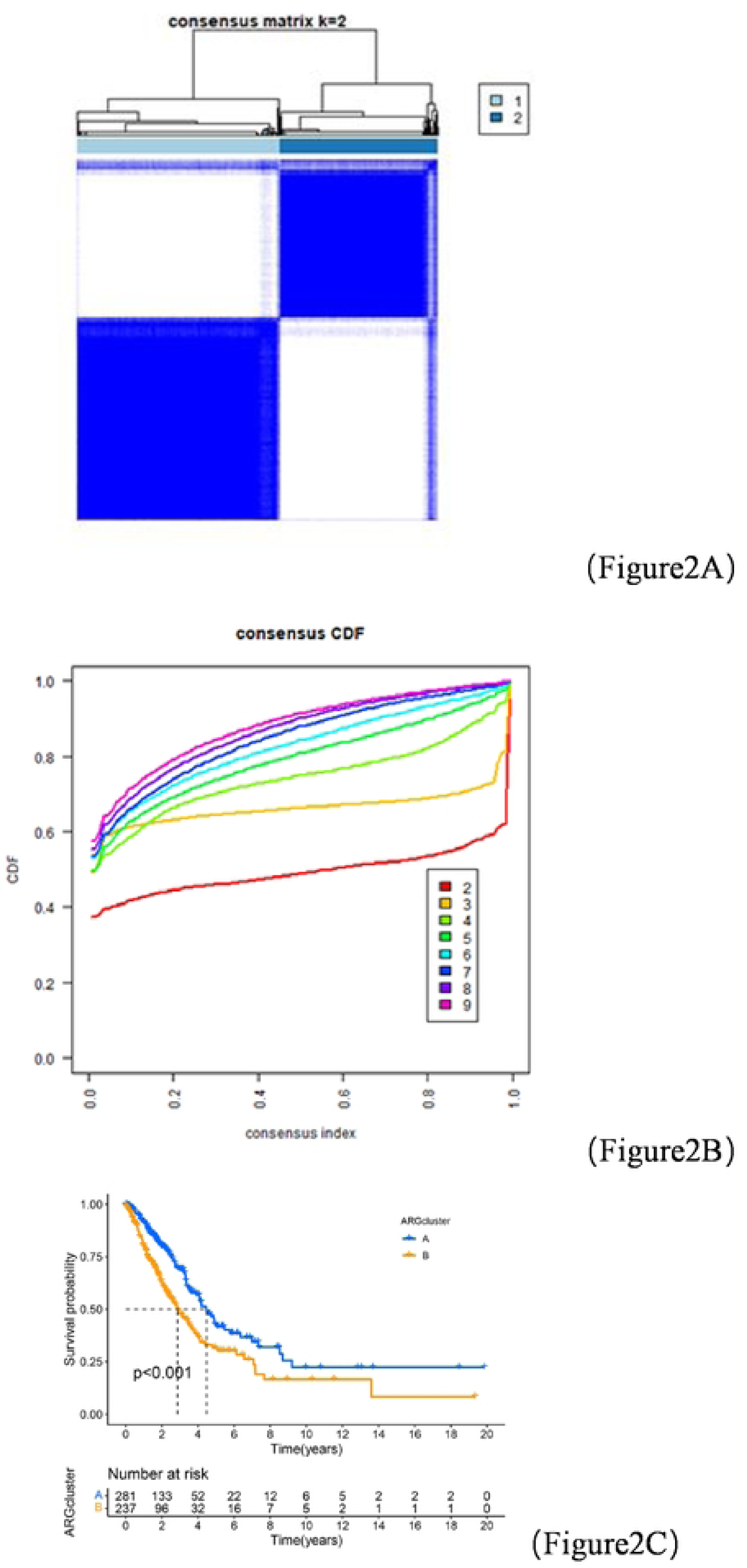
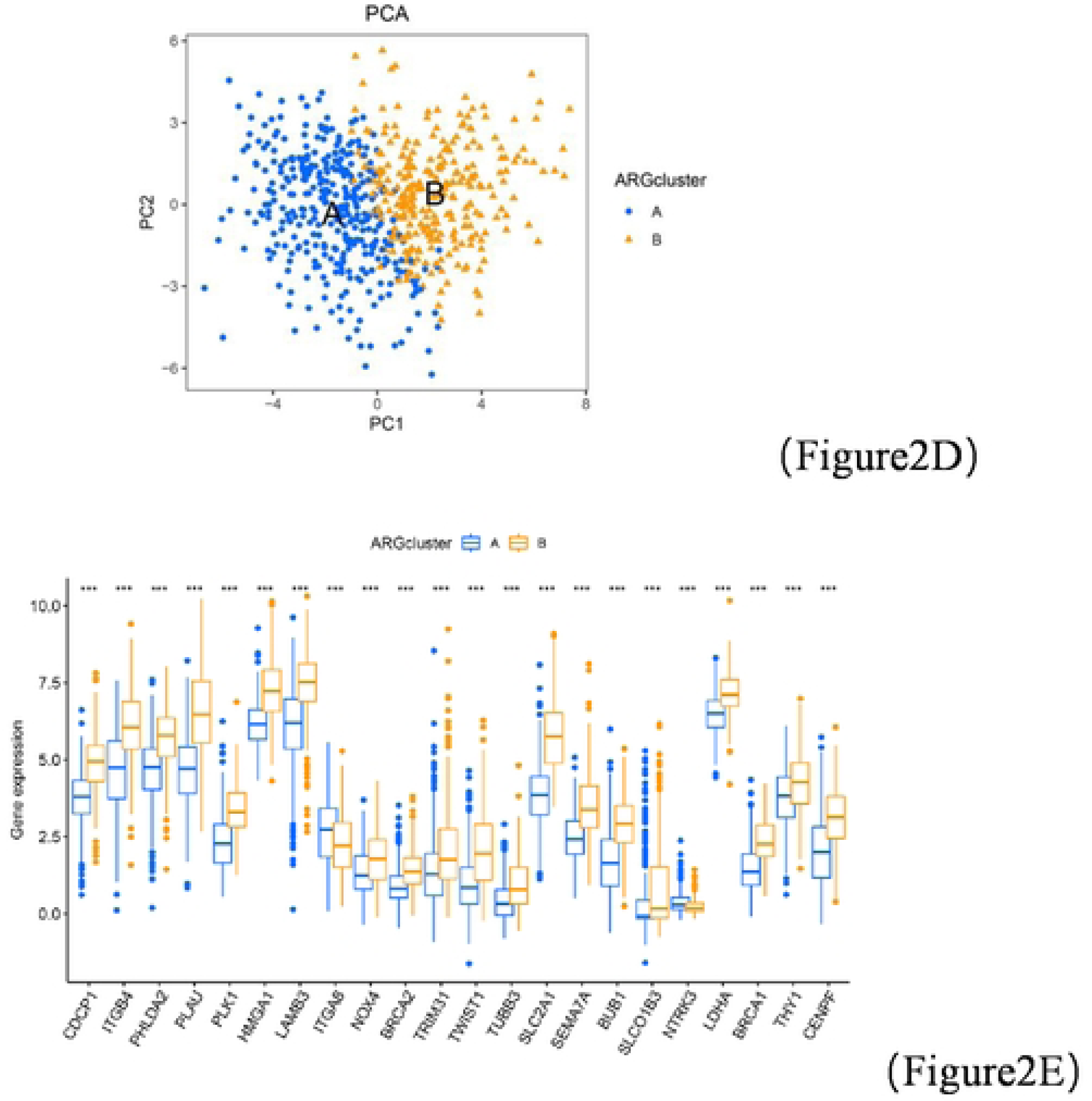
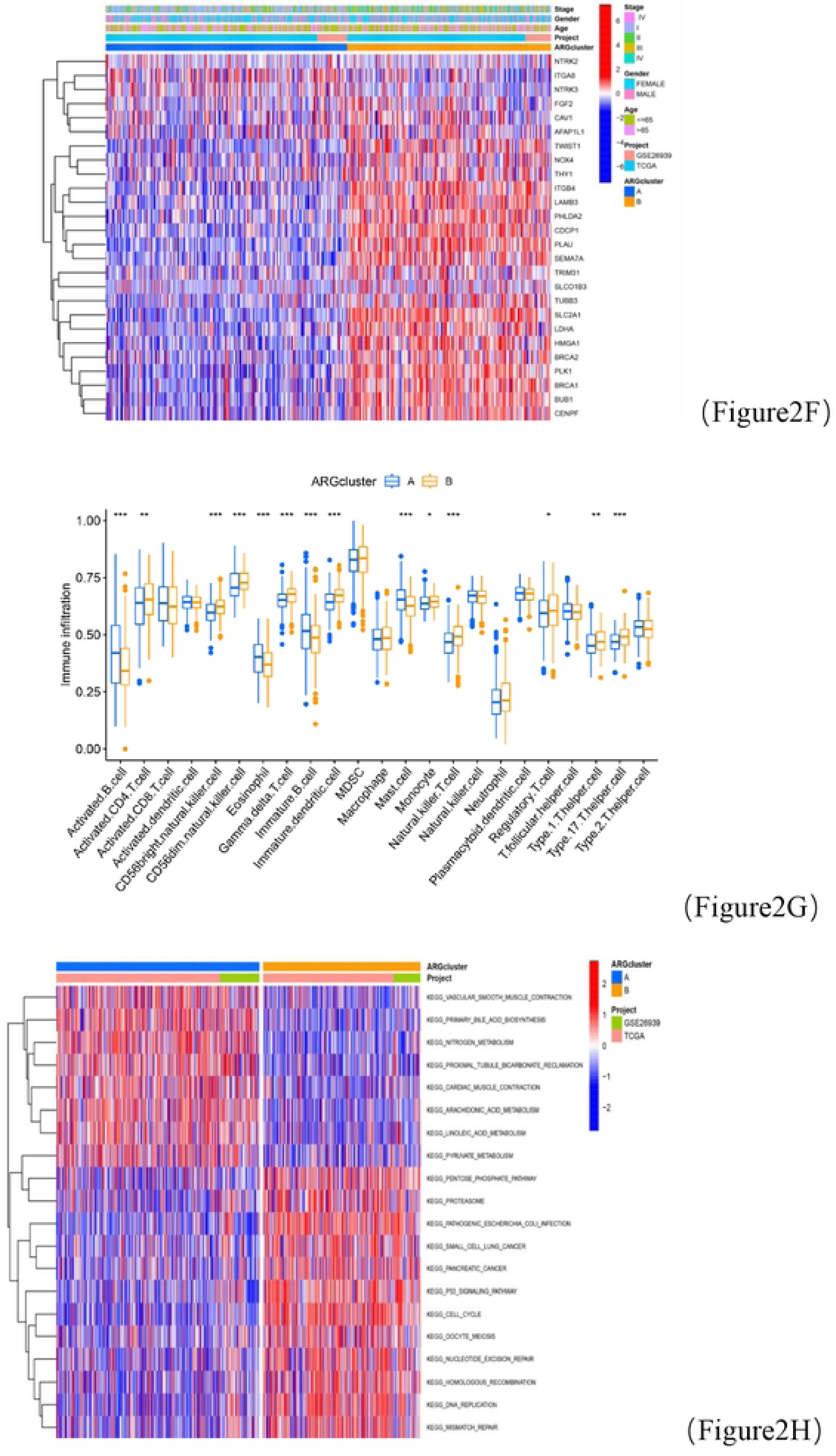
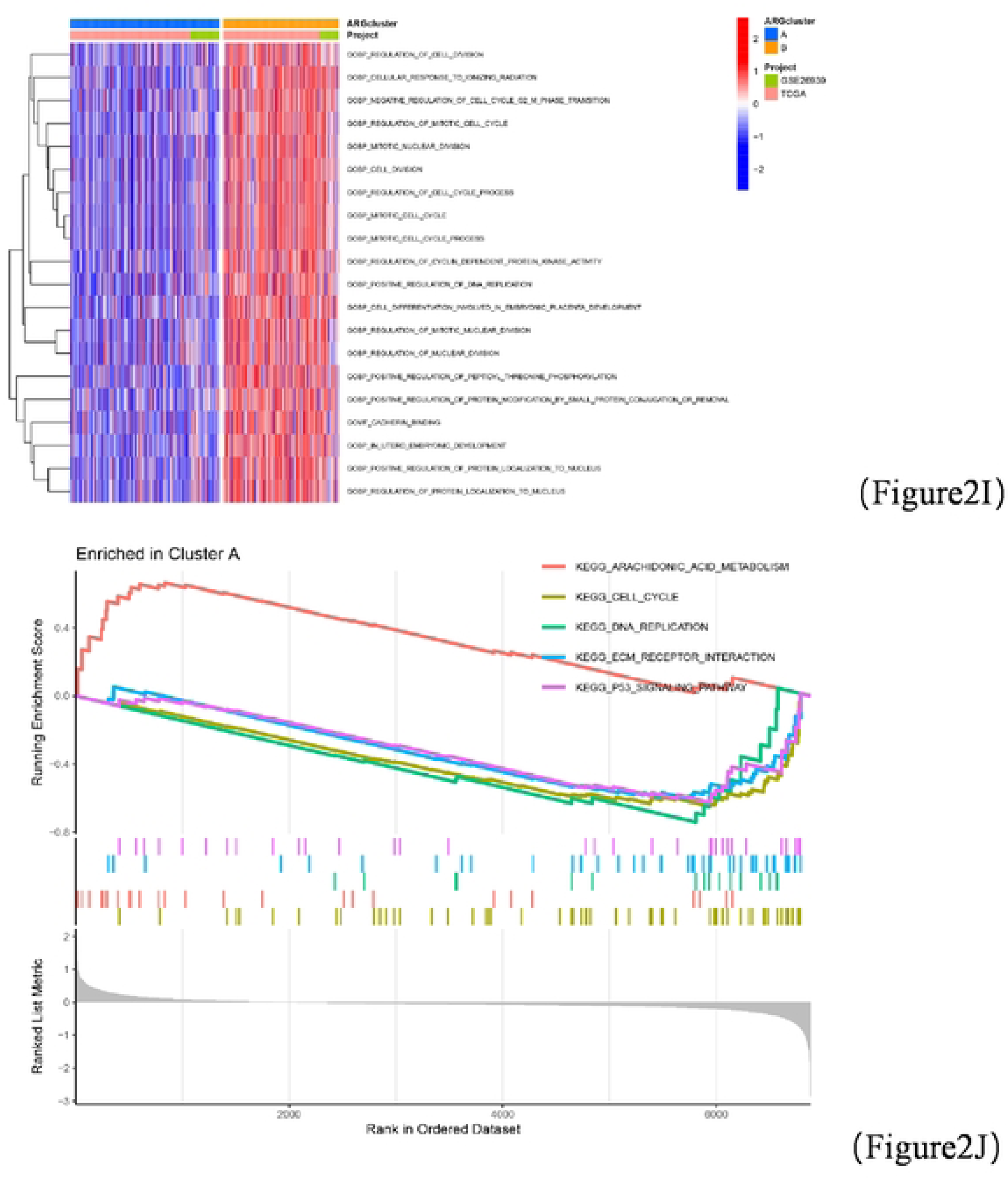
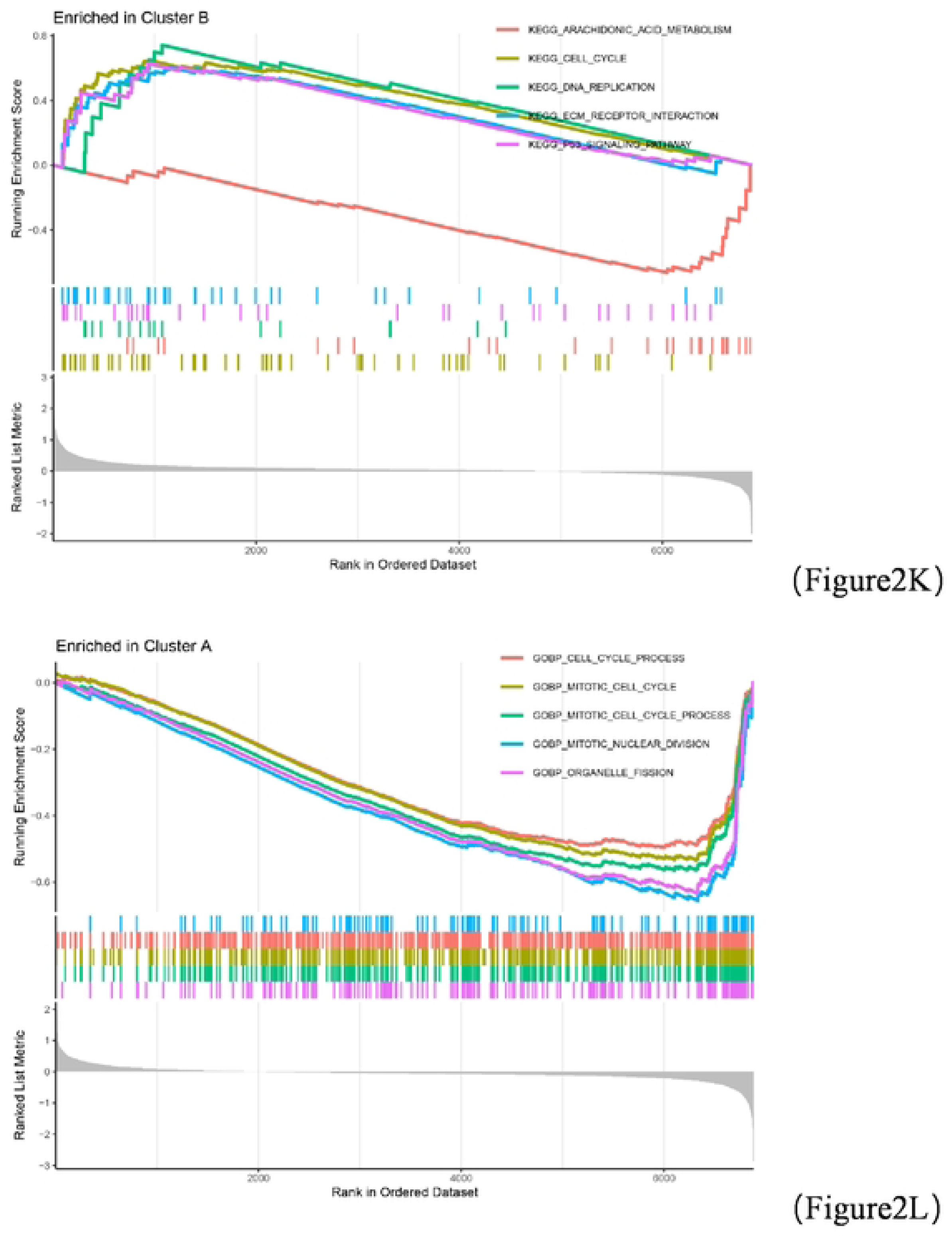
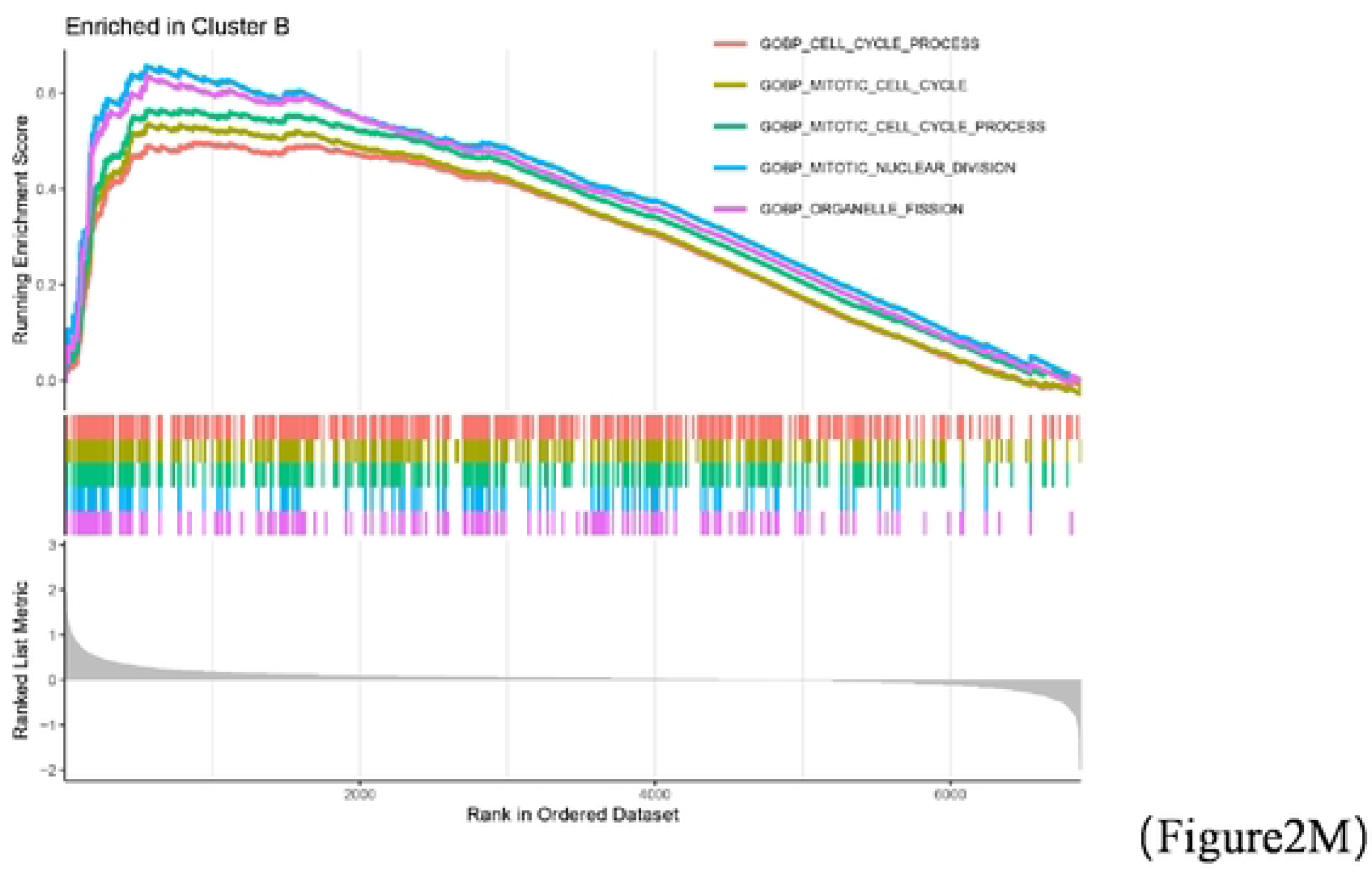
Consensus clustering analysis of Anoikis and enrichment analysis. (A-B).The results of the consensus cluster analysis analysis and the determination of the optimal K value. (C). Analysis of KM curves between clusters (D). PCA analysis. (E). Expression of differential genes between subgroups. (F). Relationship between Anoikis genes and clinical variables. (G). Immune cell infiltration levels between different clusters. (H-I). Functional enrichment analysis of GO, KEGG between different clusters. (J-M). ssGSVA analysis between subgroups(p < 0.05 *; p < 0.01 **; p < 0.001 ***)

### Construction of prognostic model

The included samples were randomly divided into an train cohort (260) and a test cohort (260). Previously, for the initial 26 genes obtained that were associated with prognosis in lung adenocarcinoma patients, we attempted to use LASSO and multifactorial analysis(Figure 3A-B). Five gene sets NTRK2, TRIM31, SLCO1B3, LDHA, THY1 were finally identified. Finally, a risk score was constructed using five genes associated with anoikis that predicted patient survival and prognosis., and the ARG_score was accessed as described: Risk score =ARG score=NTRK2*-0.276250142994865+TRIM31*0.128590889689903+SLCO1B3*0.188473392753616+LDHA*0.3507142 31730417+THY1*0.334794624752464. Next, we take the median value of the obtained risk scores and classify the test cohort and the whole cohort into high and low risk groups based on the median value. K-M analysis showed a significant difference in OS between the high first risk groups in the training set (Figure 3C). We also validated the accuracy of ROC to predict the prognosis of GBM patients (Figure 3D). Also, the OS of the test cohort and the whole cohort were compared. The results are similarly different(Figure3E-F). We also validated the accuracy of ROC in predicting prognosis of LUAD patients in a training cohort (Figure 3F). The predicted 1-year, 3-year, and 5-year survival rates were respectively 0.777, 0.654 and 0.661. Also, the test cohort and the whole cohort were validated(Figure 3G-H). Multifactorial Cox regression analysis was used to assess whether clinical characteristics (gender, age, stage) and risk score were independent prognostic factors for the patients. We found that advanced stage and risk score were independent prognostic factors for patients (Figure 3I, Table S13). As shown in Figure 3J, the expression of the selected genes was increased in the high-risk group, except for NTRK2.The distribution of patients between groups with different grouping patterns (Figure 3K). Meanwhile, we further analyzed the risk scores between clusterA and B. We concluded that the risk score in cluster B was significantly greater than that in group A (Figure 3L).

**Figure 3.**
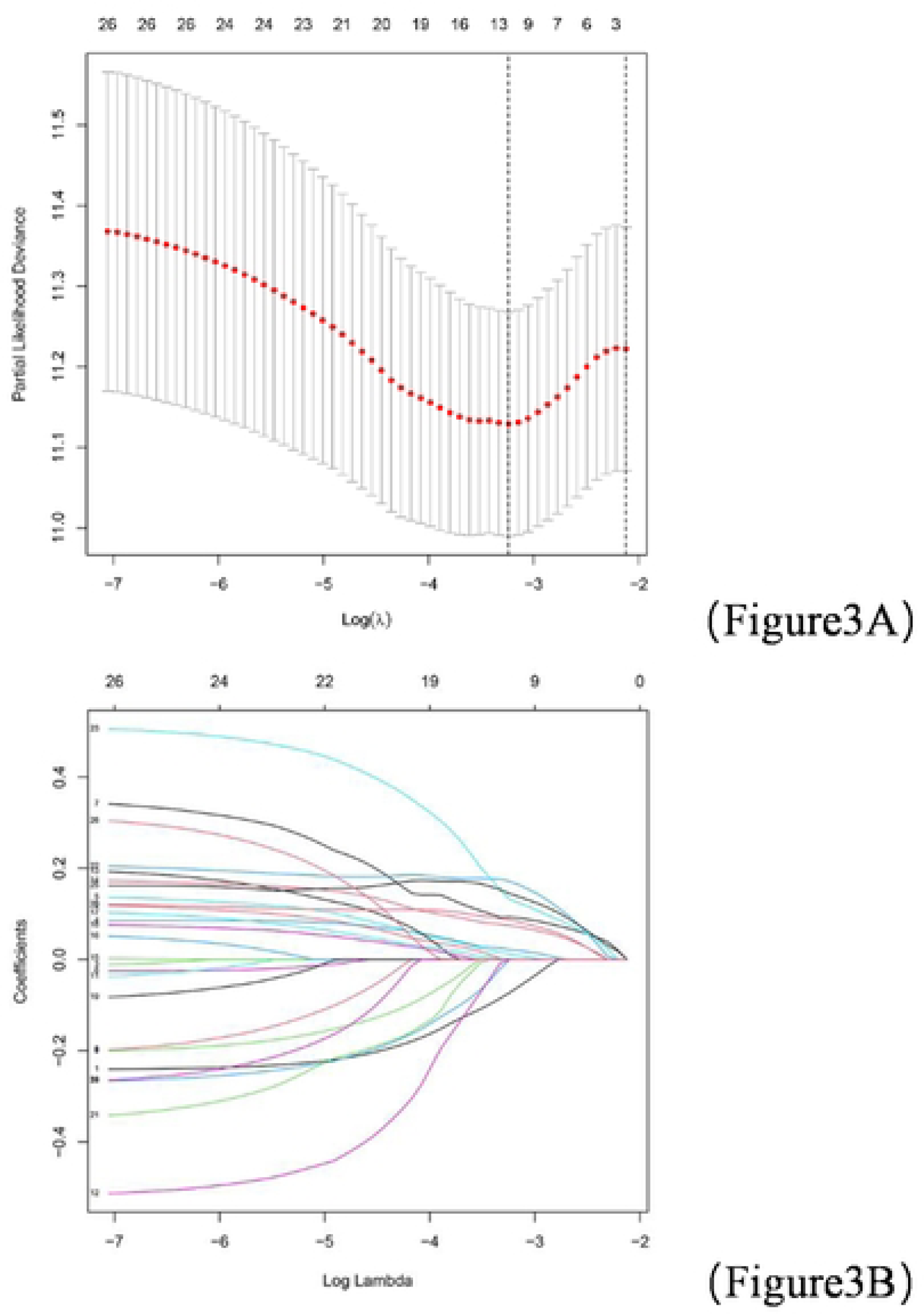
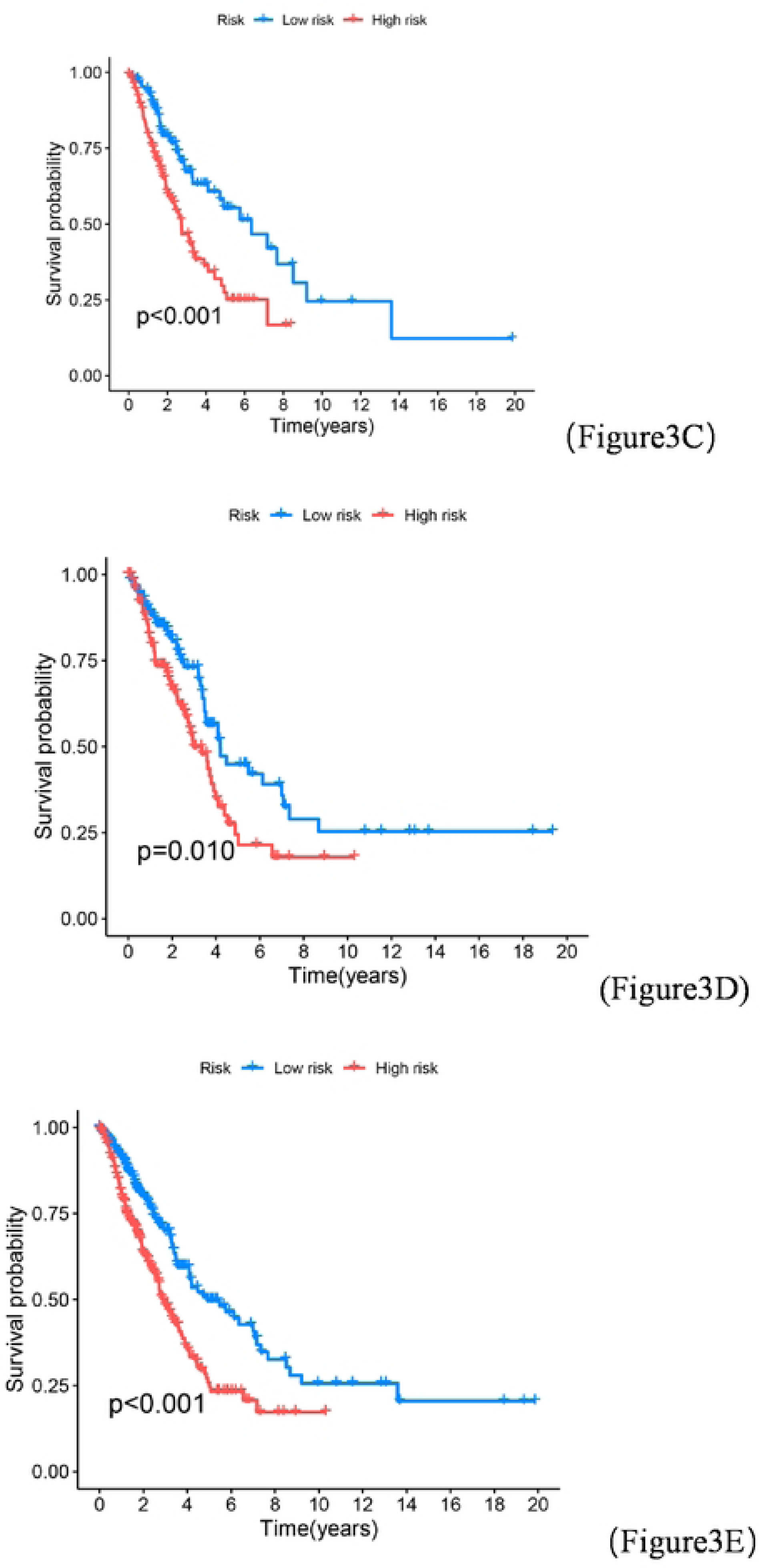
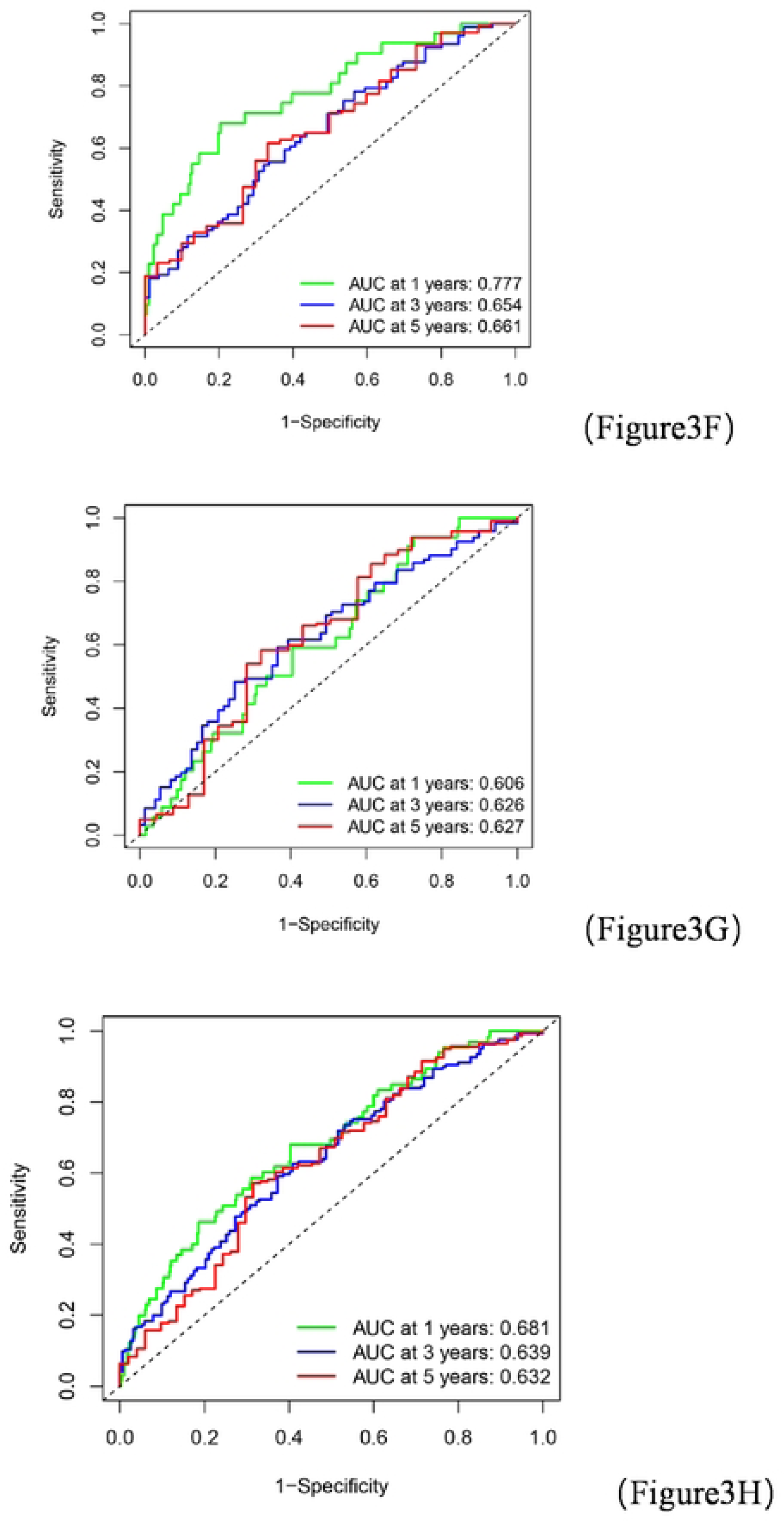
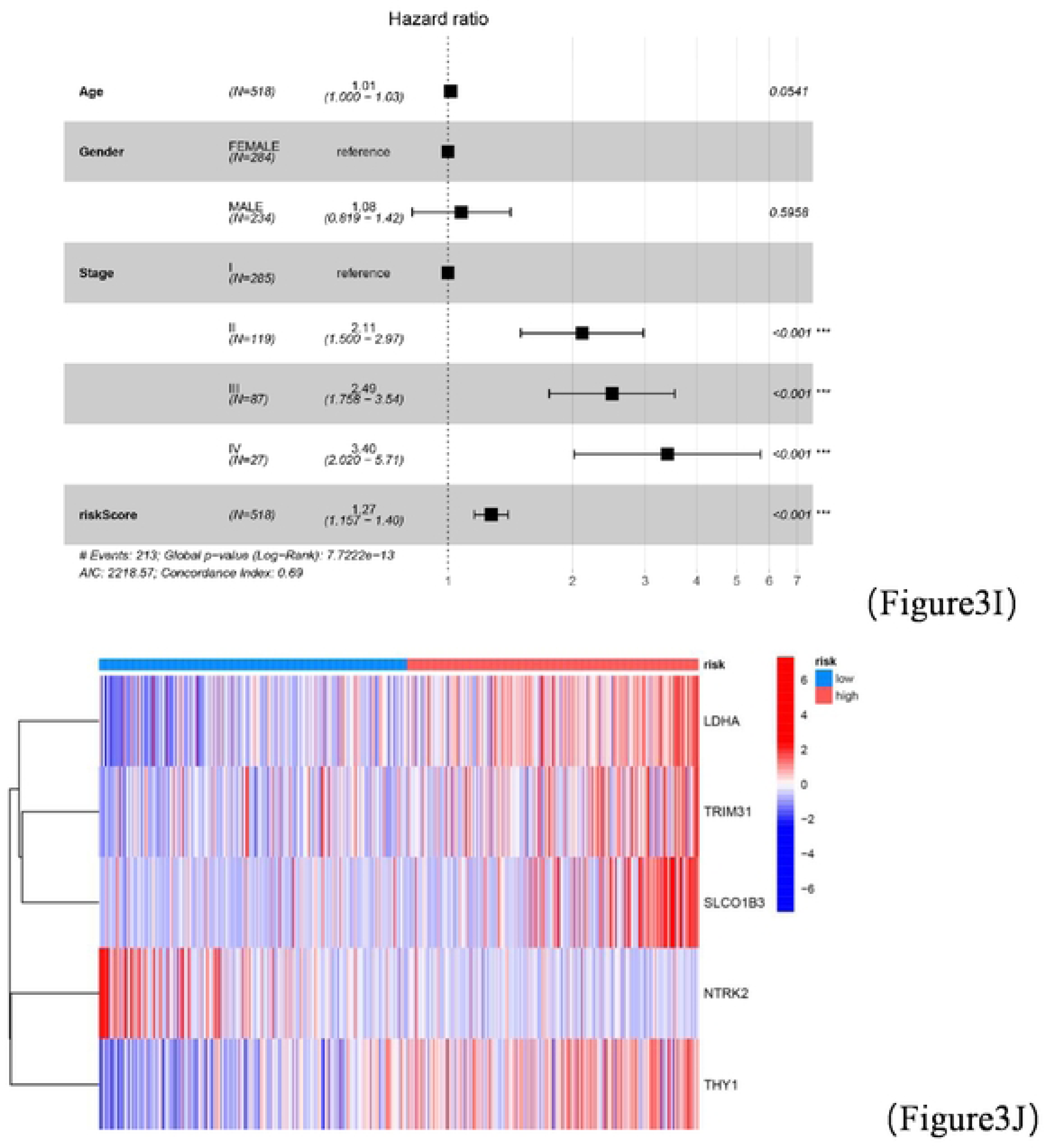
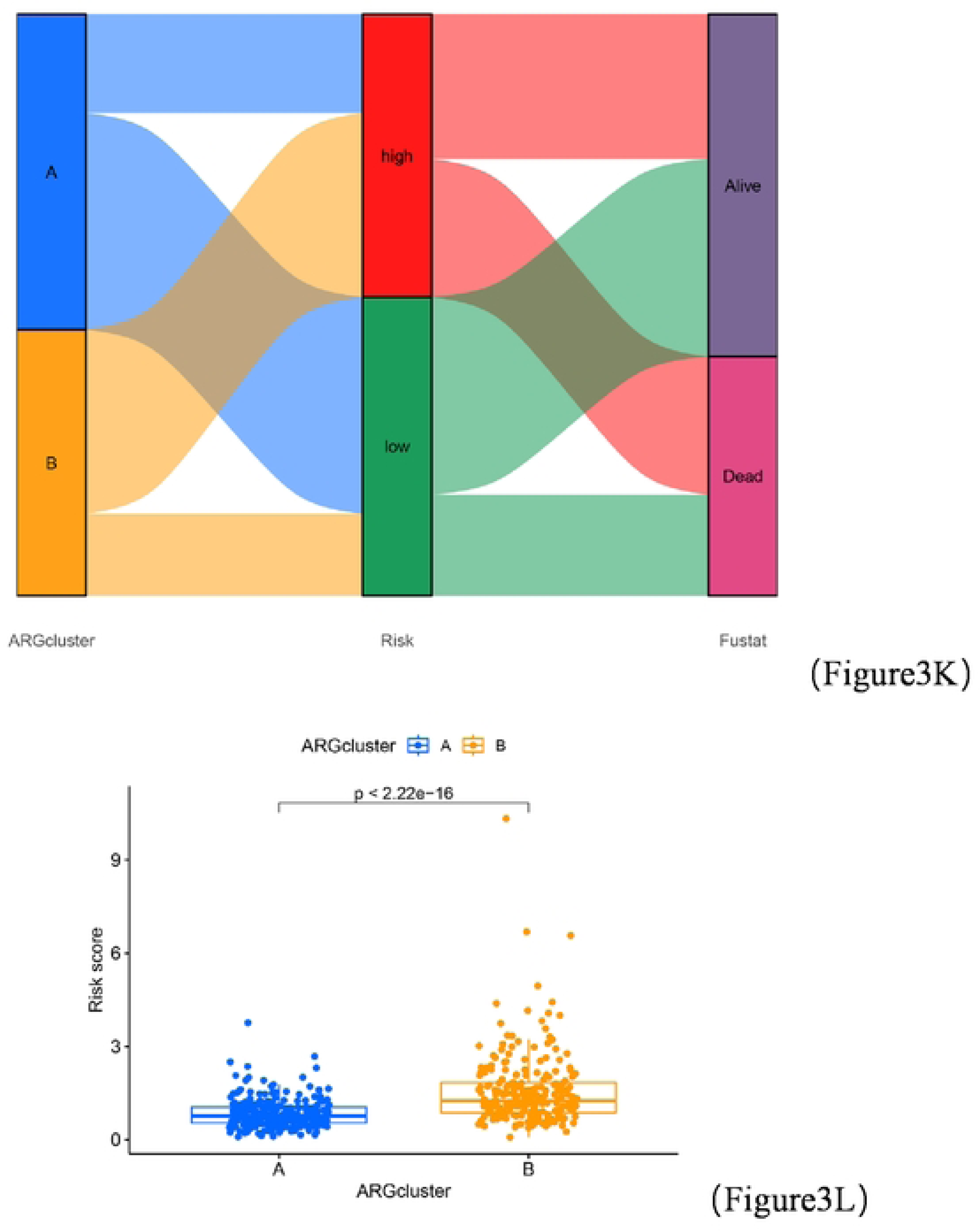
Risk score establishment. (A-B) The least absolute shrinkage and selection operator (LASSO) method of anoikis-related genes associated with prognosis. (C-E) KM curves for the training cohort, test cohort, and the whole cohort. (F-H) ROC curves of the training, test, and the whole cohort. (I) Forest plots of the results of multifactorial analysis. (J)Risk heat map of the selected genes. (K) Alluvial plot of the distribution of samples among different groups. (L) Comparison of risk scores between apoptotic gene subgroups(p < 0.05 *; p < 0.01 **; p < 0.001 ***).

### A nomogram constructed based on risk scores and clinical variables

We constructed a nomogram that predicts the probability of survival at 1, 3 and 5 years for patients with LA based on high and low risk (Figure 4A). The calibration curves confirmed the high predictive accuracy of our constructed Nomogram (Figure 4B). From the cumulative risk curves, it is clear that the risk tends to increase with increasing years in both the high and low risk groups, while the risk is significantly higher in the high risk group than in the low risk group (Figure 4C). This prognostic model with different clinical factors has more net benefit in predicting prognosis by DCA curves (Figure 4D-F).

**Figure 4.**
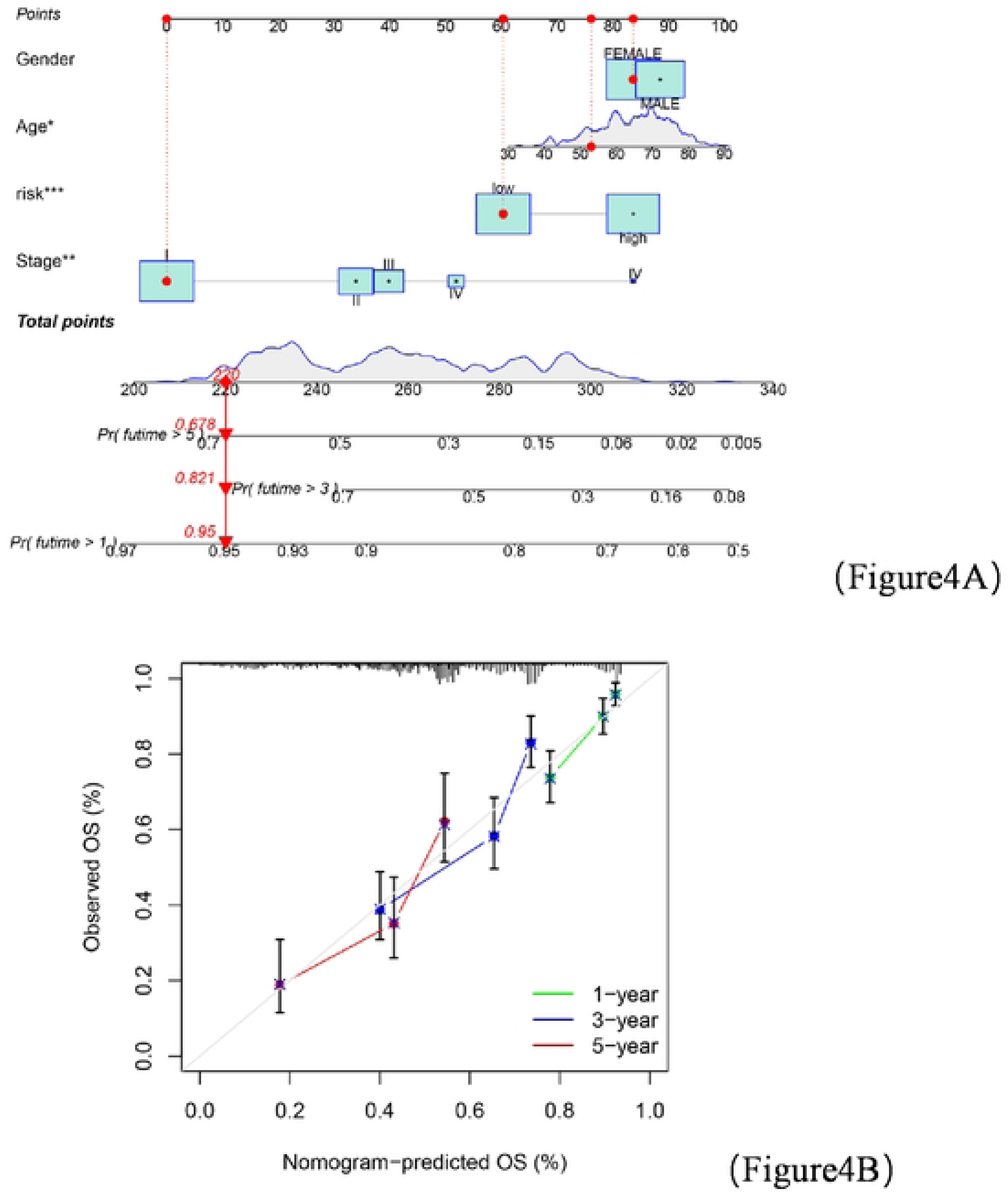
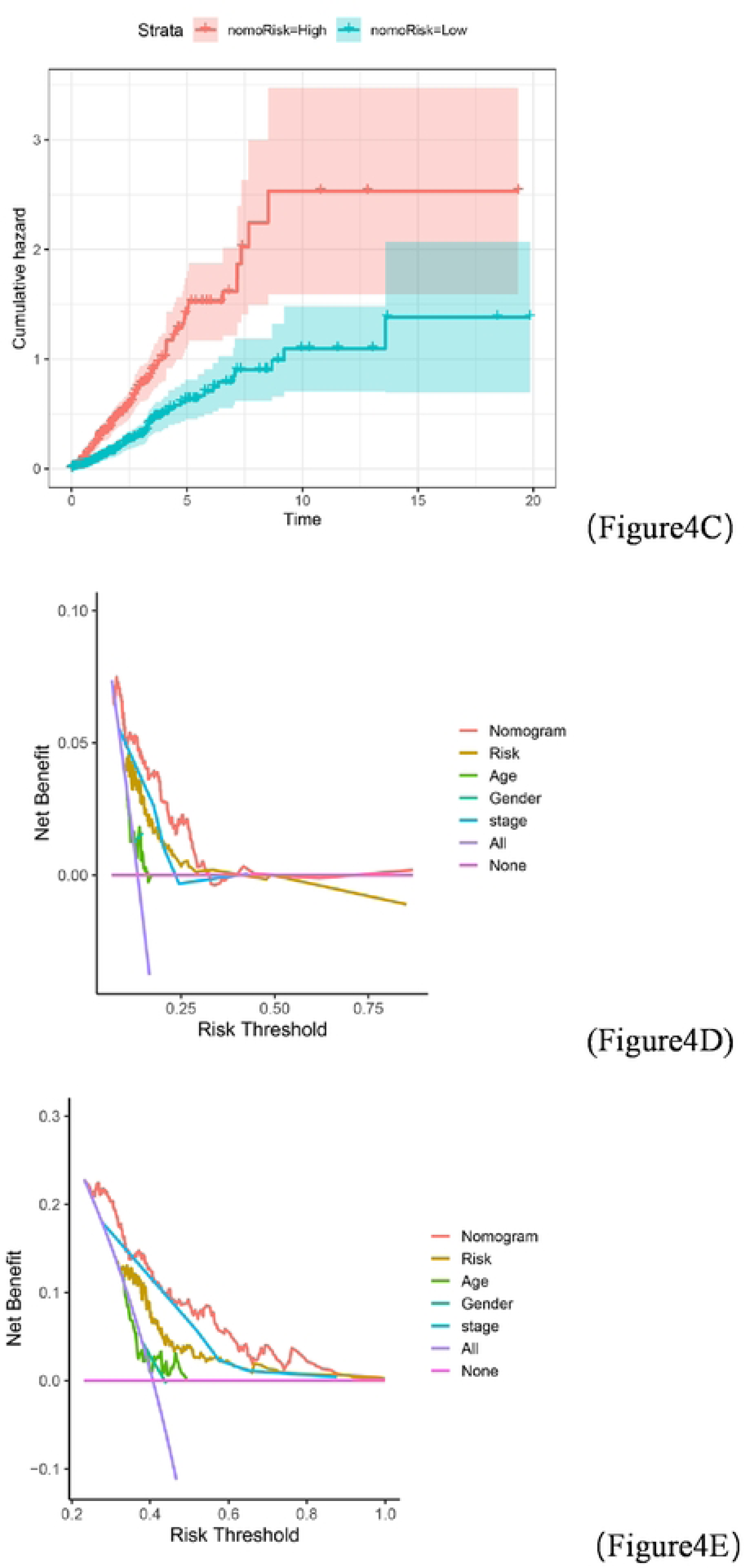
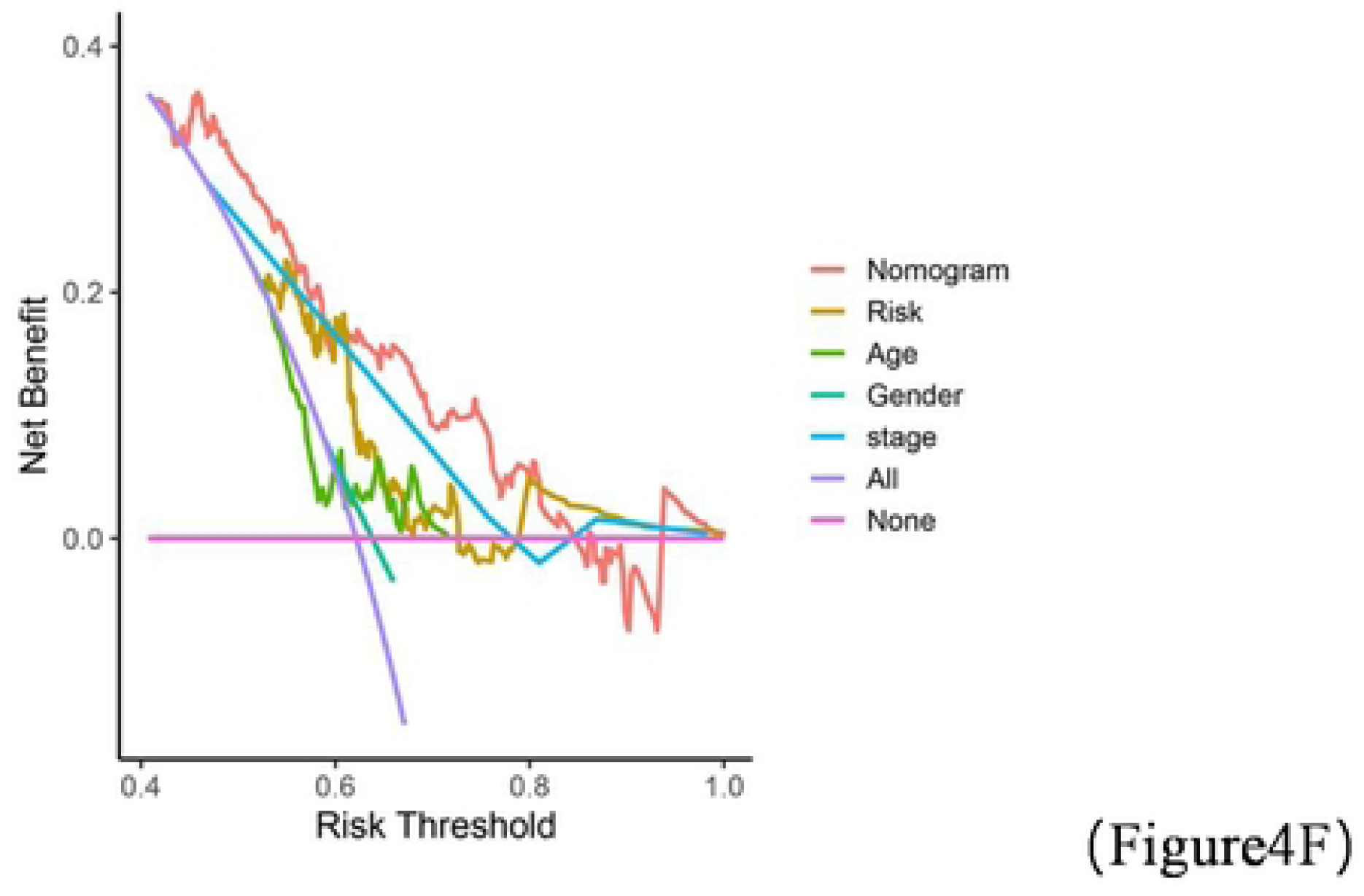
Predictive power of the risk score-based nomogram. (A) Risk score combined with clinical variables constructed by nomogram. (B) Calibration curves. (C) Cumulative risk curves between high and low risk groups. (D-F) DCA curves comparing 1-, 3-, and 5-year OS of the nomogram, respectively (p < 0.05 *; p < 0.01 **; p < 0.001 ***).

### Immune status between different risk groups, TME analysis

As shown in Figure 5A, we performed correlation analysis between immune cells, and it can be seen from the figure that most of the immune cells have positive or negative regulatory relationships with each other. We then compared the levels of immune cells between the different risk groups. Figure 5B shows the percentage of immune subtype cells between the different groups. In addition to this, we compared the immune subtype cell content between the two groups and found that a few cells (DC cells, macrophagesM1, neutrophils,B cell memory, mast cell activated) differed between the risk groups (Figure 5C, TabLe S14).The relationship between each subtype of cells and risk score was also evaluated (as in Figure 5D). The relationship between immune subtype cells and risk scores can be clearly shown to be positive between Macrophages M0, mast cell activated, neutrophils, T cells CD4 memory activated, NK cell resting and risk scores, while the rest are negatively correlated. And, we evaluated the correlation between selected genes, riskscore and immune cells.As seen in Figure 5E, there was a significant relationship between most of the genes, riskscoer and immune cells.Because the tumor microenvironment has a profound effect on the treatment outcome, we further calculated the stromal and immune scores.There was a significant difference between high and low risk groups between stromal and total scores (Figure 5F, Table S15).

**Figure 5.**
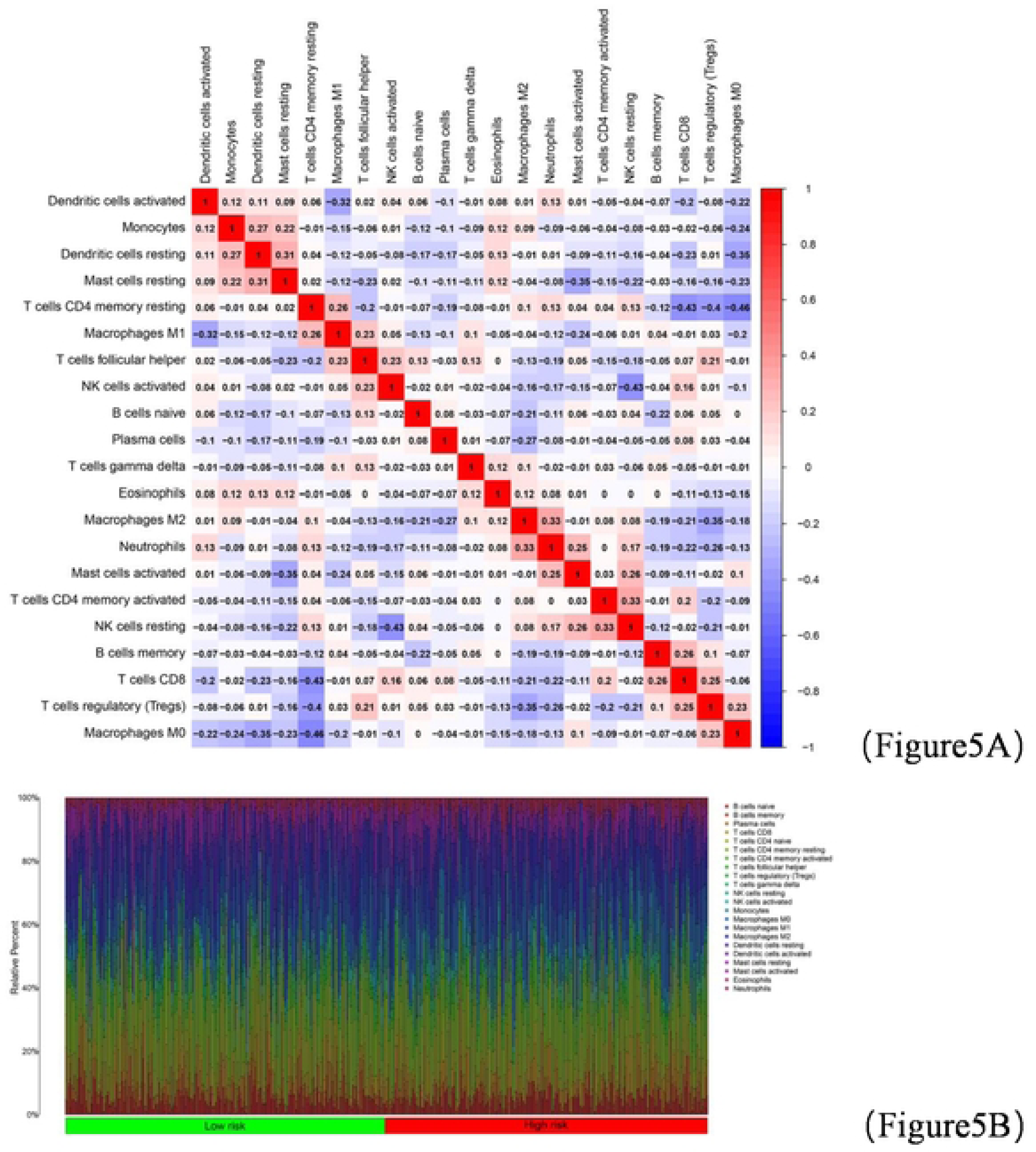
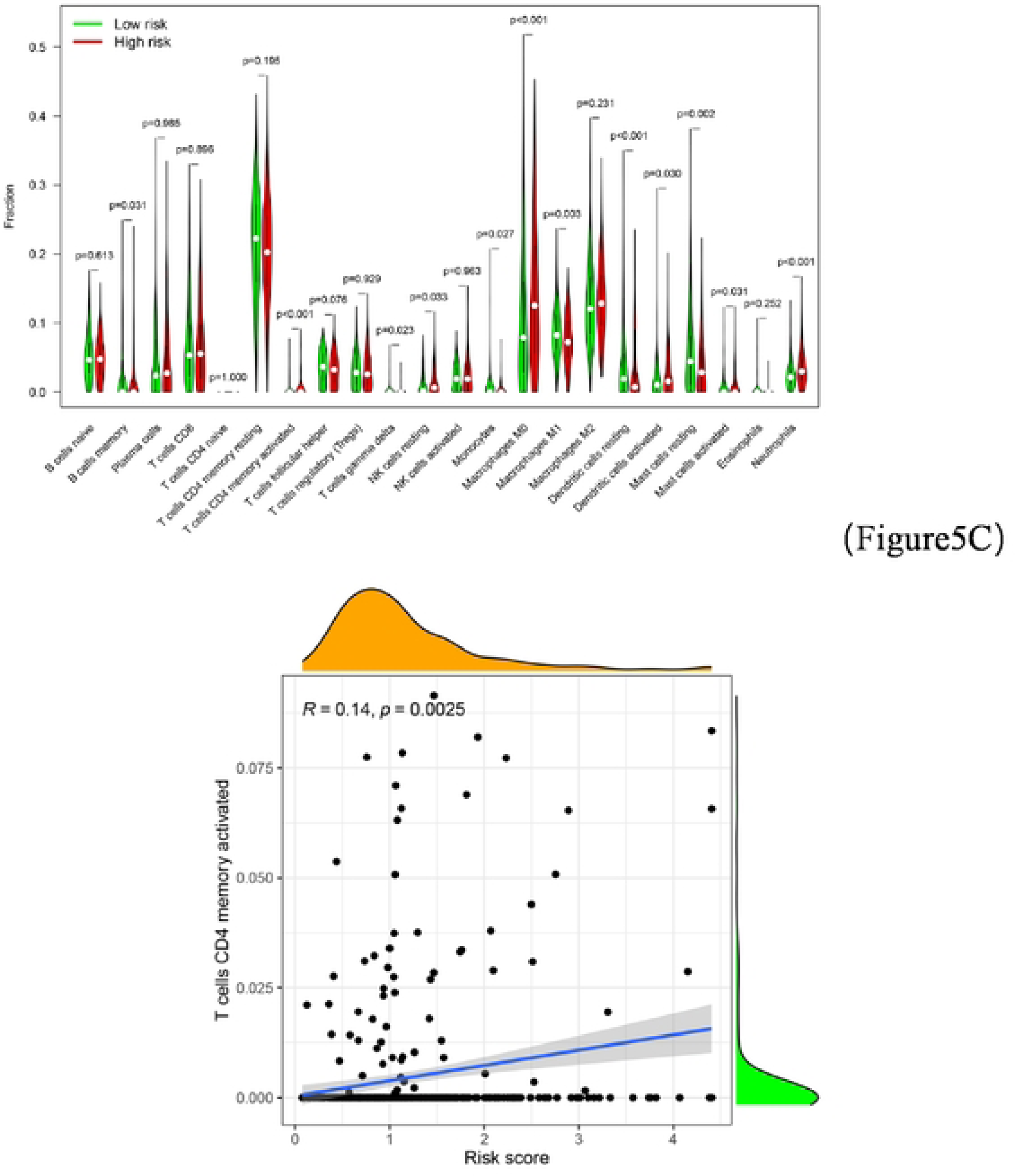
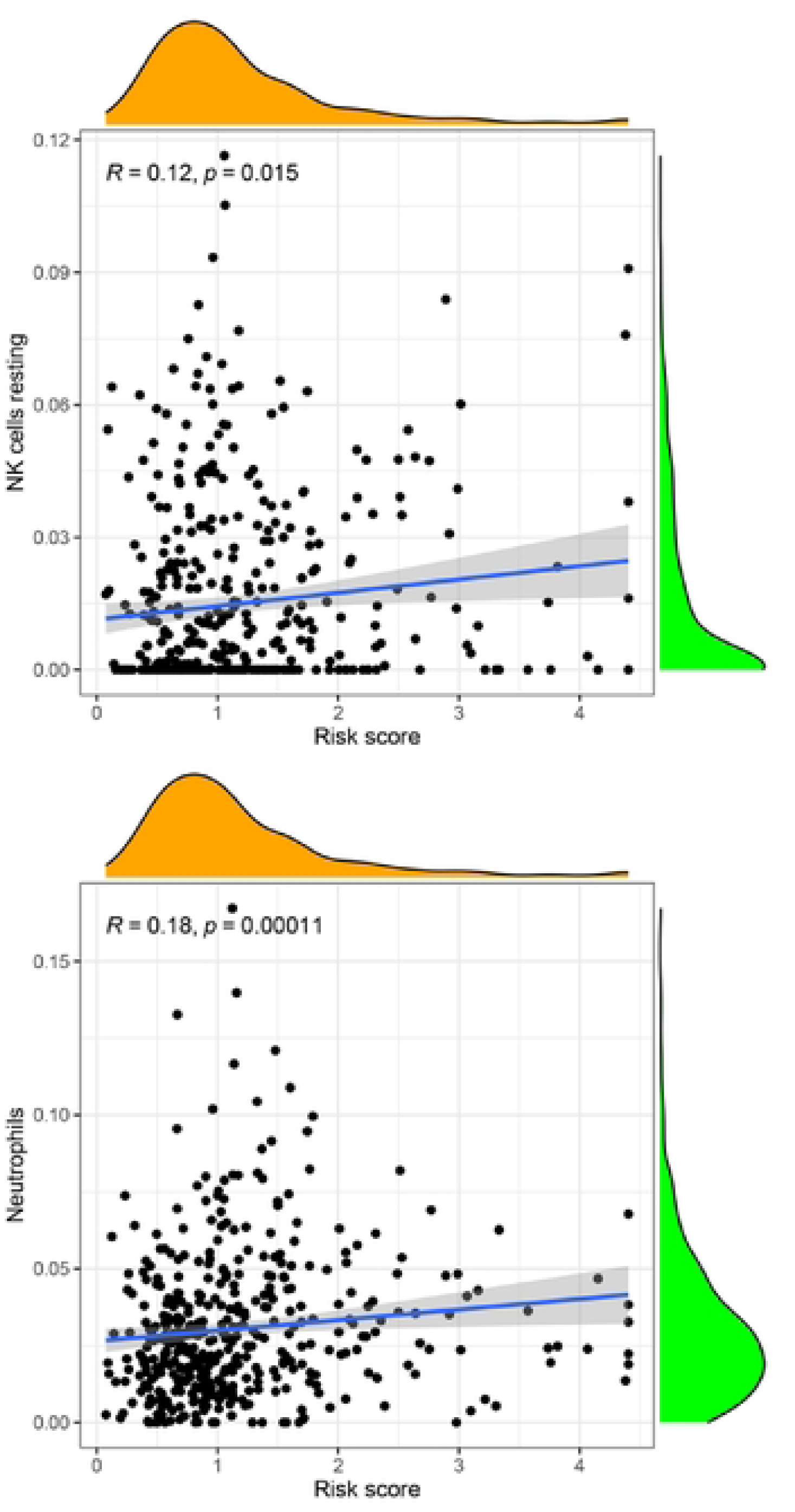
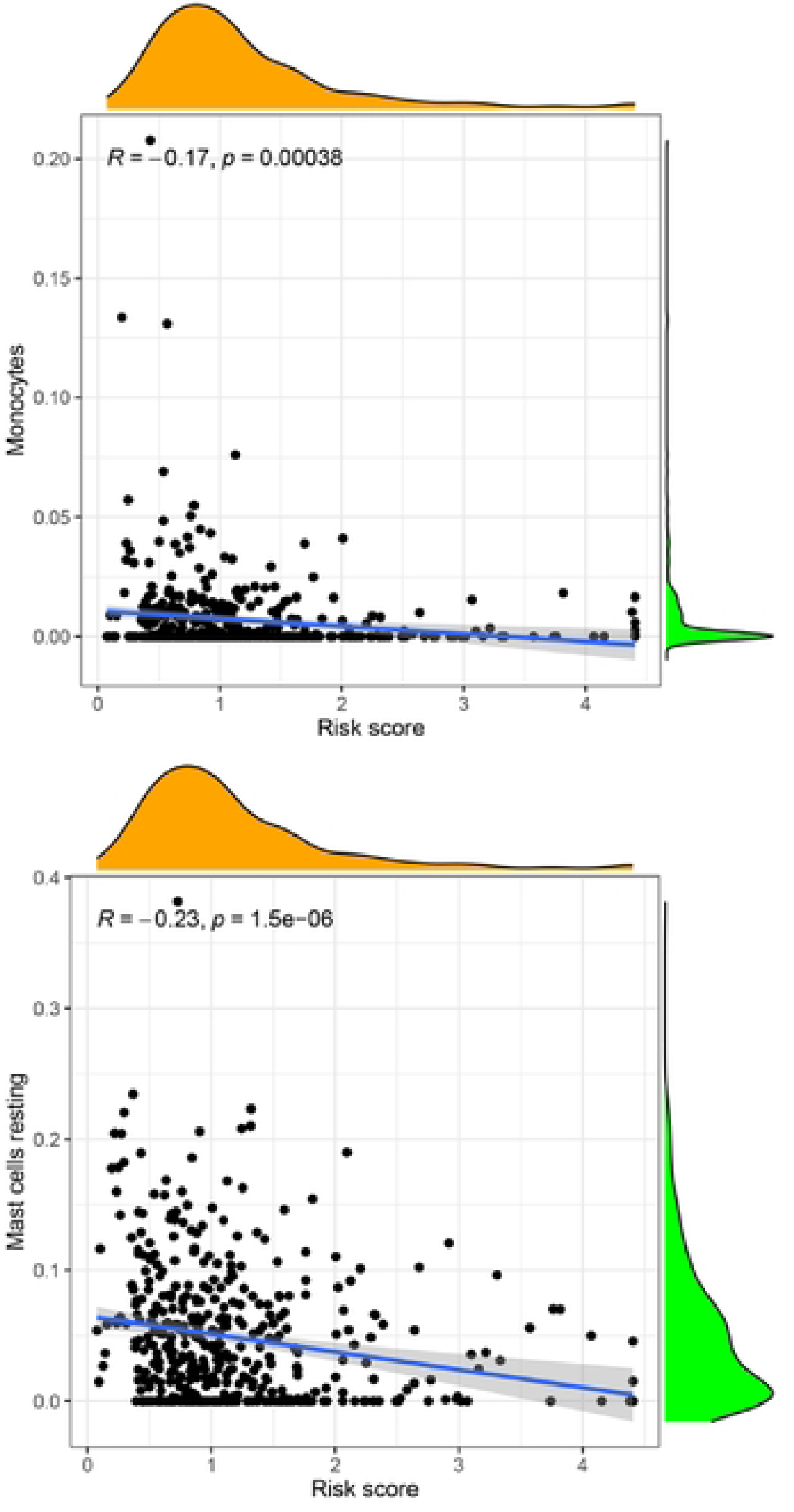
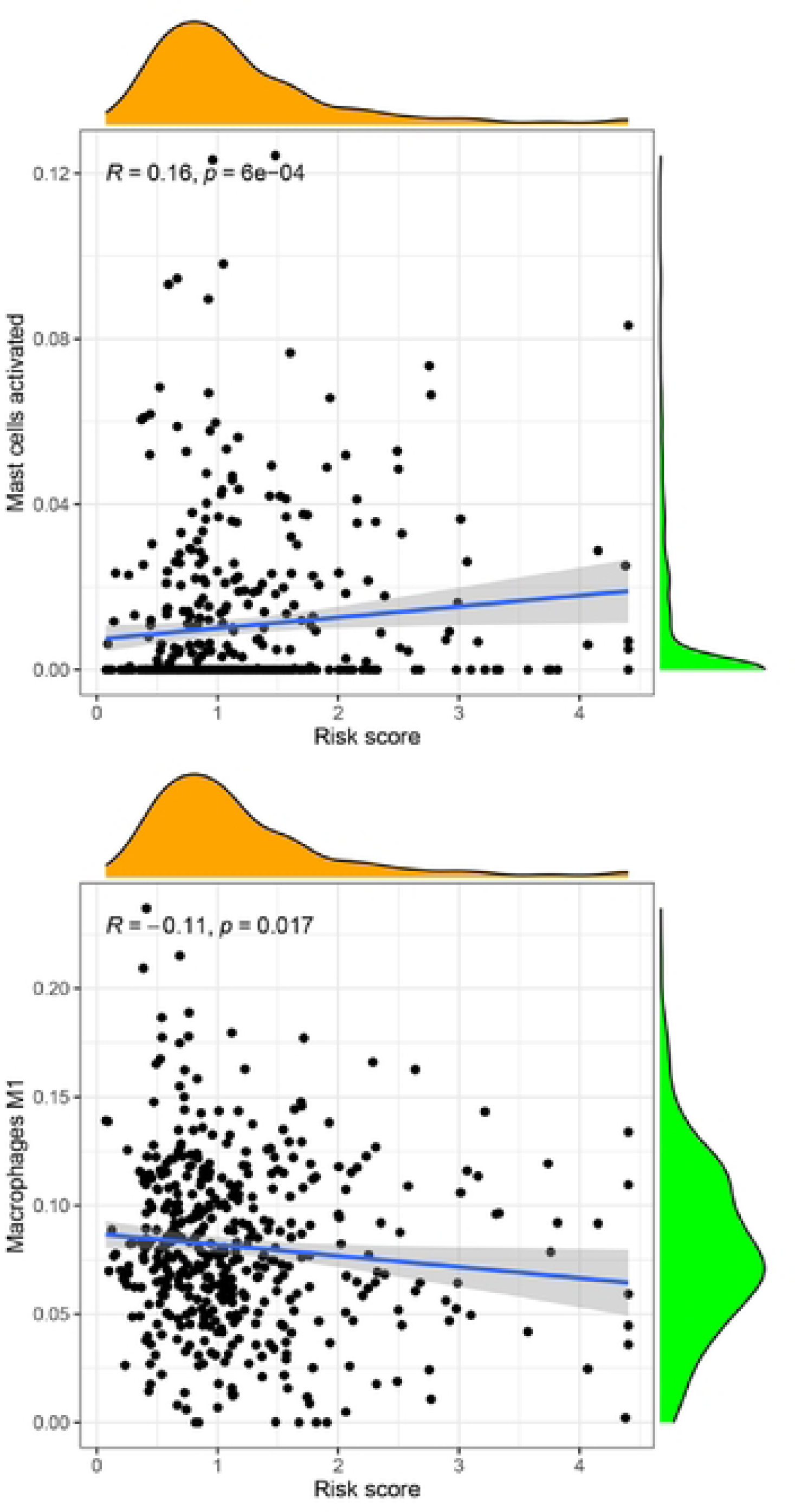
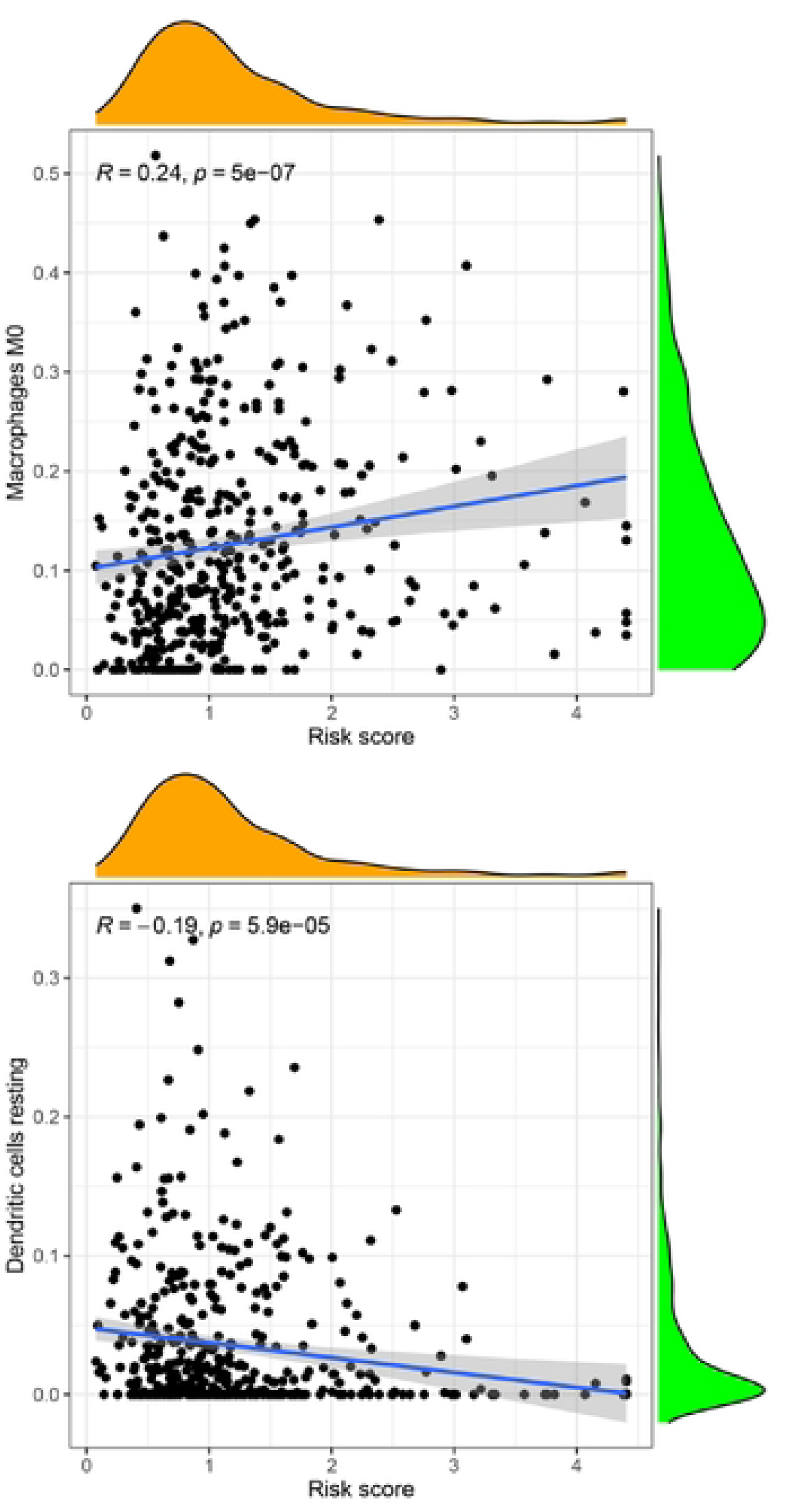
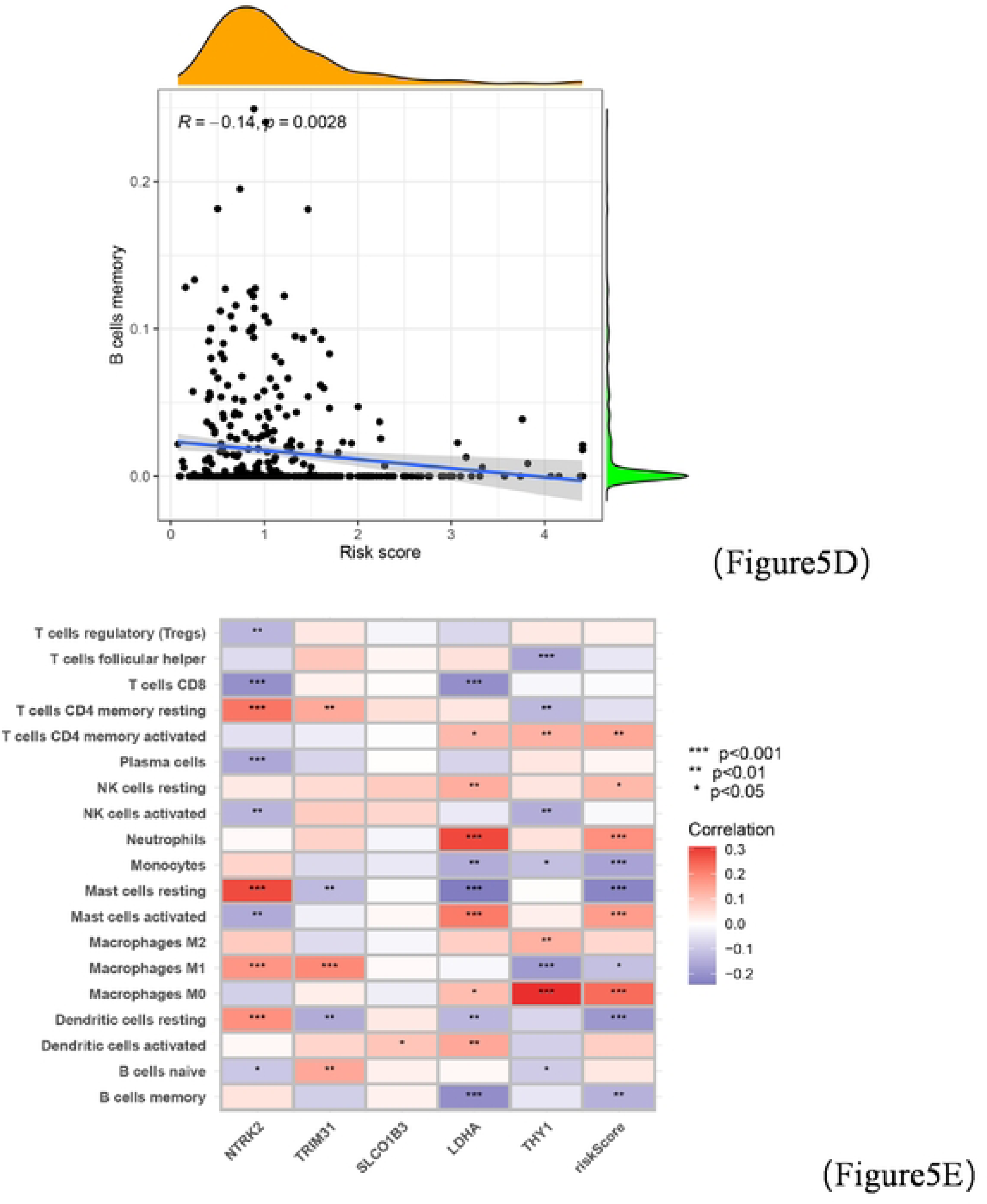
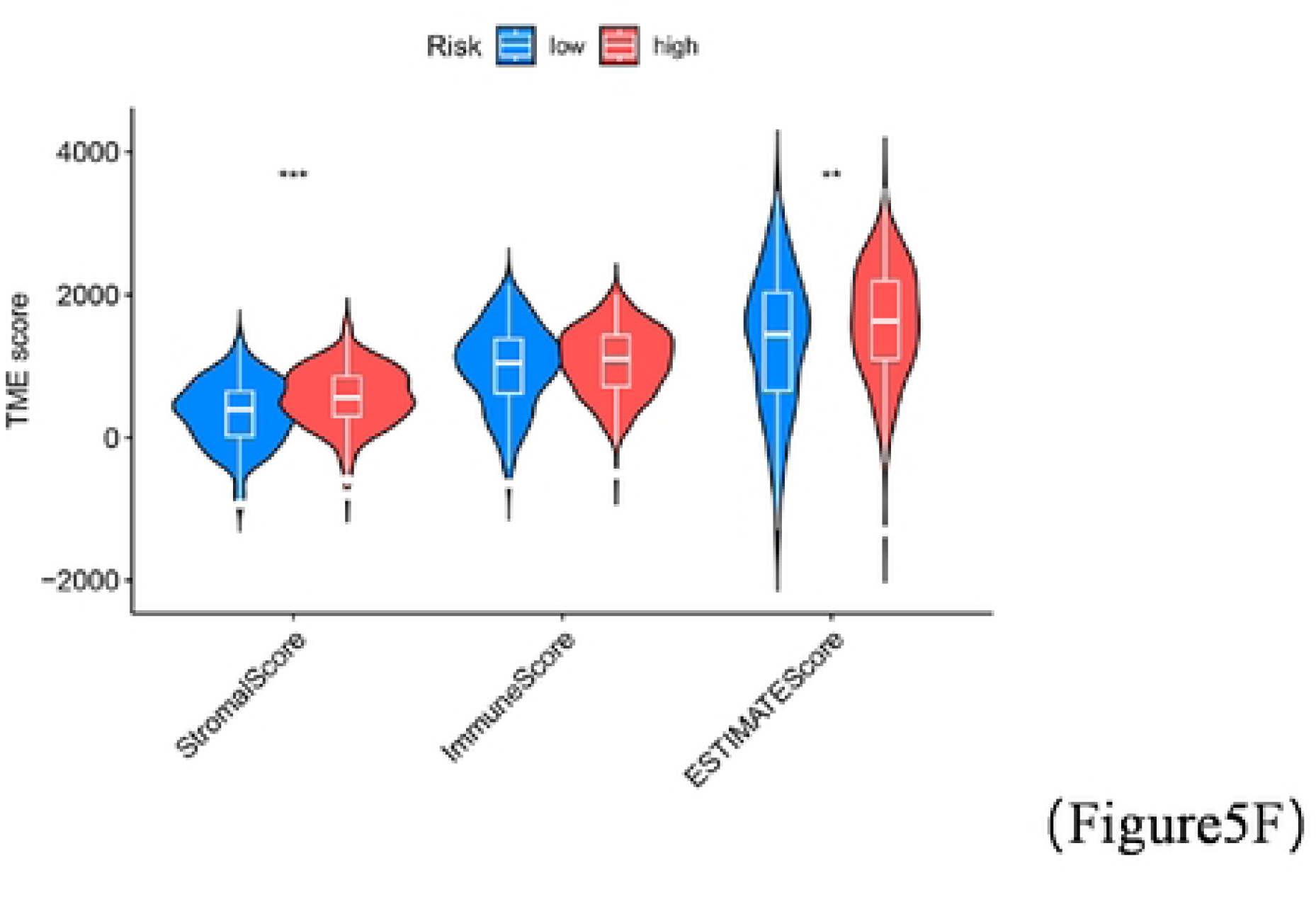
Comparison between risk score and immune status. (A) Relationship between immune cells (B) Proportion of immune cells in different samples (C) Violin plot of the abundance of immune cell infiltration between different samples (D) Relationship between risk score and immune cells (E) Relationship between risk score and selected genes and immune cells(F) Comparison of TME SCORE between different risk groups(p < 0.05 *; p < 0.01 **; p < 0.001 ***).

### Drug sensitivity analysis

We compared drug sensitivity, TIDE, and immune checkpoints(ICP) between the risk groups.As shown in Figure 6A-E, the IC50 of most common drugs was found to be significantly lower than that of the high-risk group, suggesting that the first-risk group was more sensitive to chemotherapy drugs.The TIDE scores between the two groups that were also compared were found to be significantly different between the scores of the high-risk group and the low-risk group, and the high-risk group was higher than the low-risk group. We concluded that immunotherapy was better in the low-risk group (Figure 6F, Table S16). Finally, we compared the ICP and found significant differences between the two groups (Figure 6G, Table S17).

**Figure 6.**
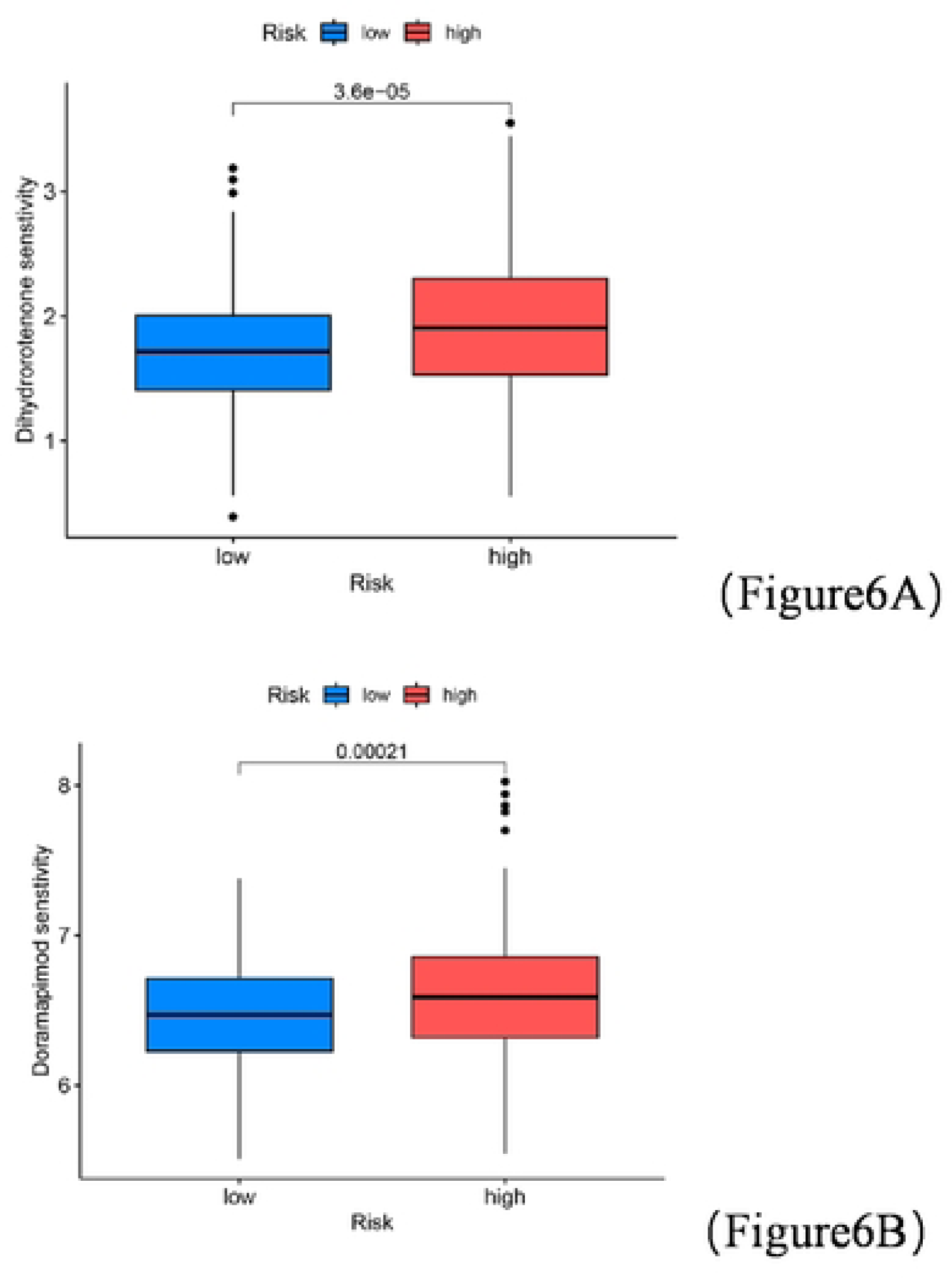
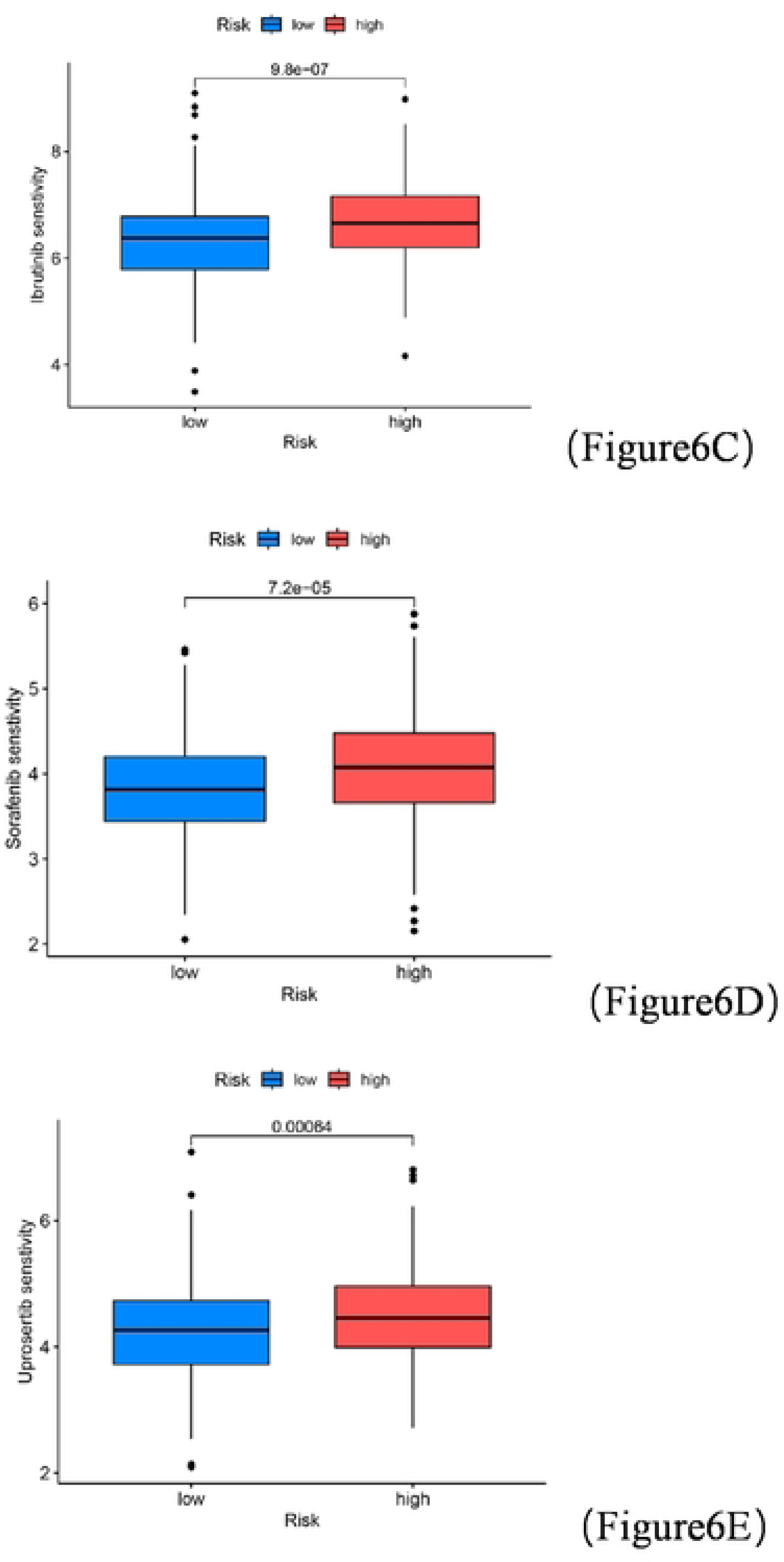
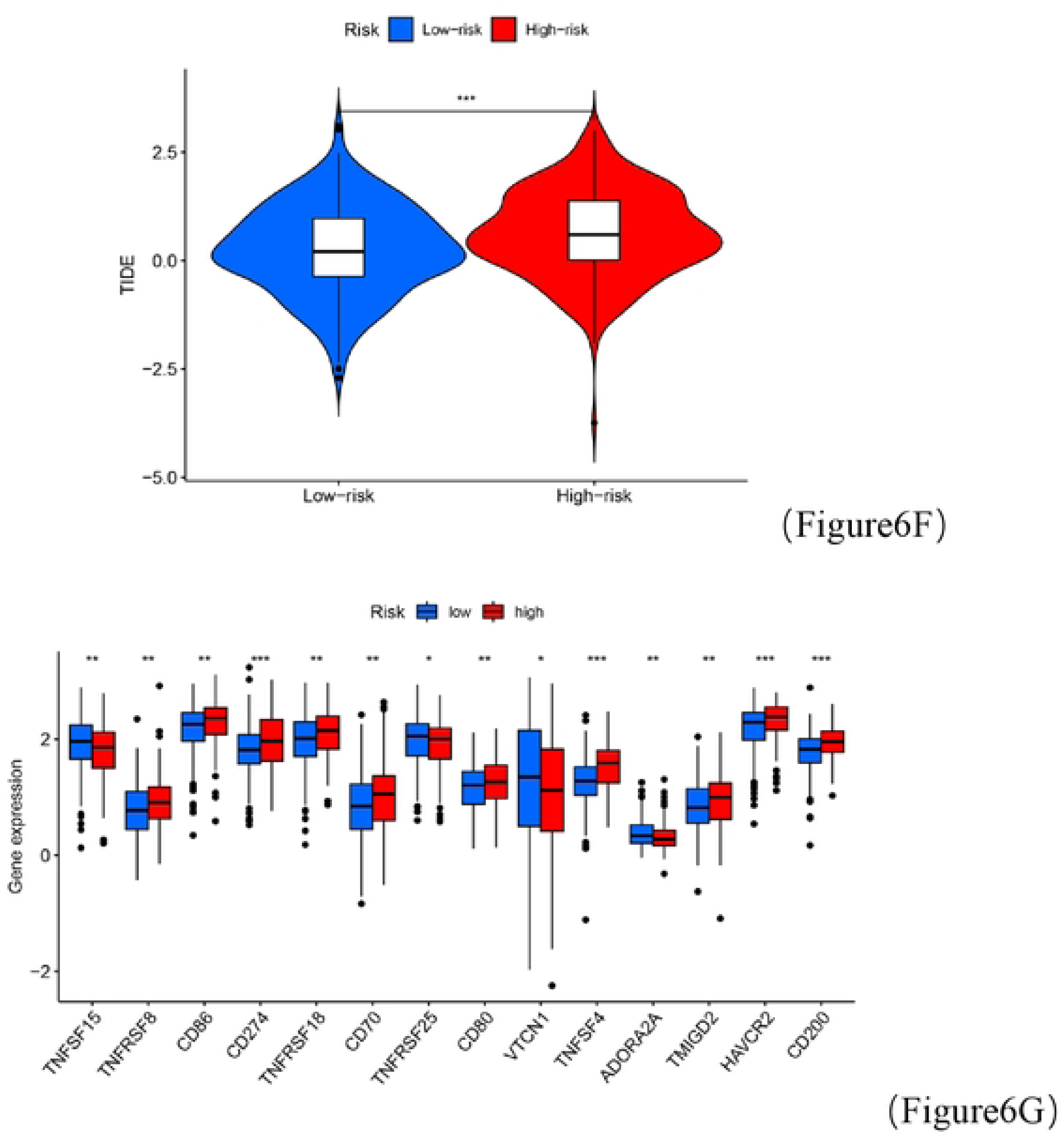
Sensitivity analysis of common drugs. (A-E) Sensitivity analysis of drugs not used between the two groups. (F) Tumor immune dysfunction and exclusion between the two groups. (G) Comparison between ICP(p < 0.05 *; p < 0.01 **; p < 0.001 ***).

### Single cell data analysis

We finally selected the complete NSCLC single cell data byTISCH2 website (https://tish.comp-genomics.org/) to find the relationship between the selected genes and cells. We selected GSE131907 for analysis, and the cell annotation results are shown in Figure 7A, 7B. Figure 7C reveals the proportion of cells in each fraction, and we can see from the figure that CD4 T cells are the most abundant, followed by CD8 T cells. Figure 7D shows the content of various immune cell subtypes in the samples. Finally, we determined the relationship between the selected genes and cell types (Figure 7E-7L). LDHA was the most abundantly expressed and the most abundantly expressed of the selected genes in DC cells.

**Figure 7.**
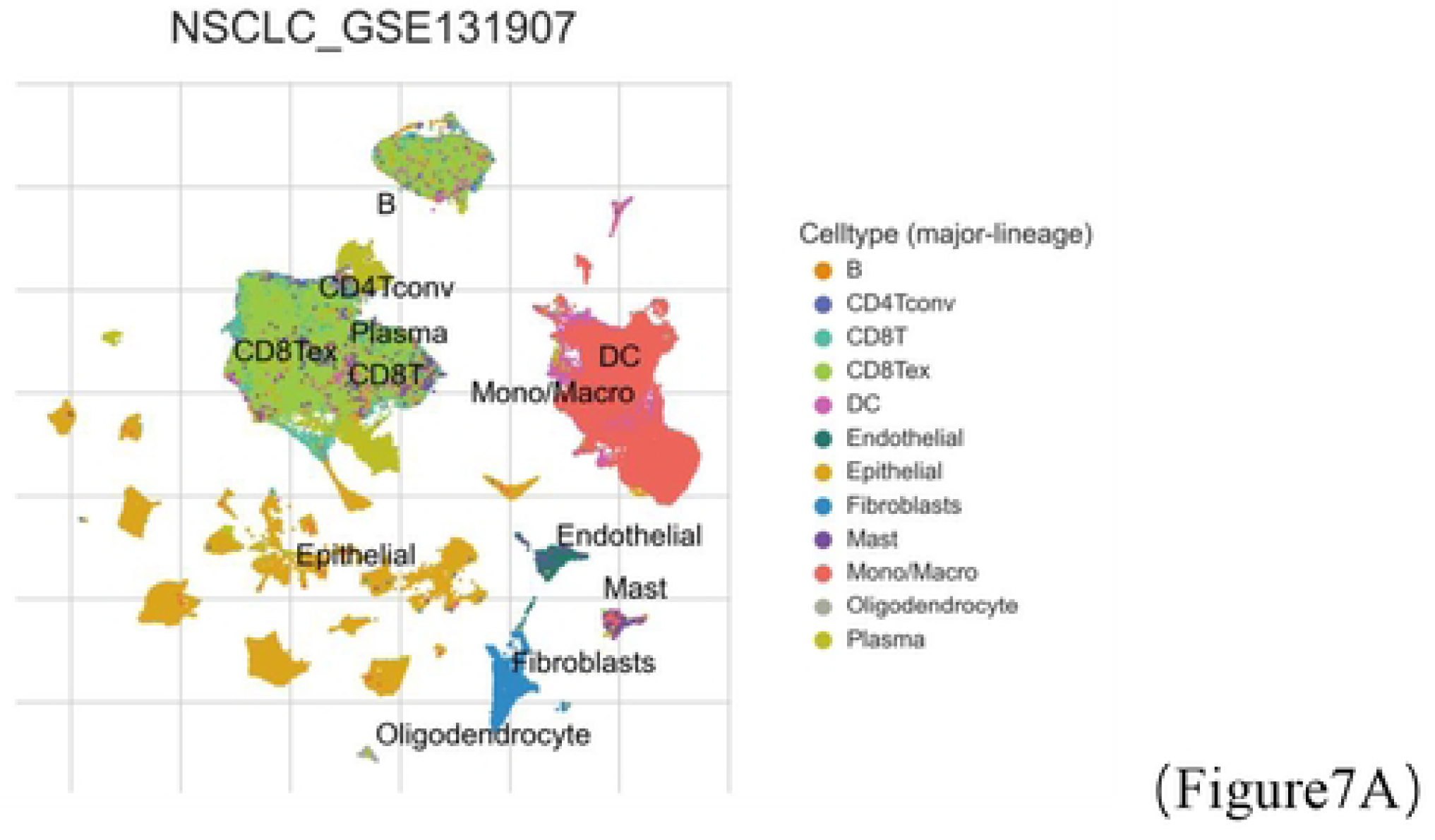
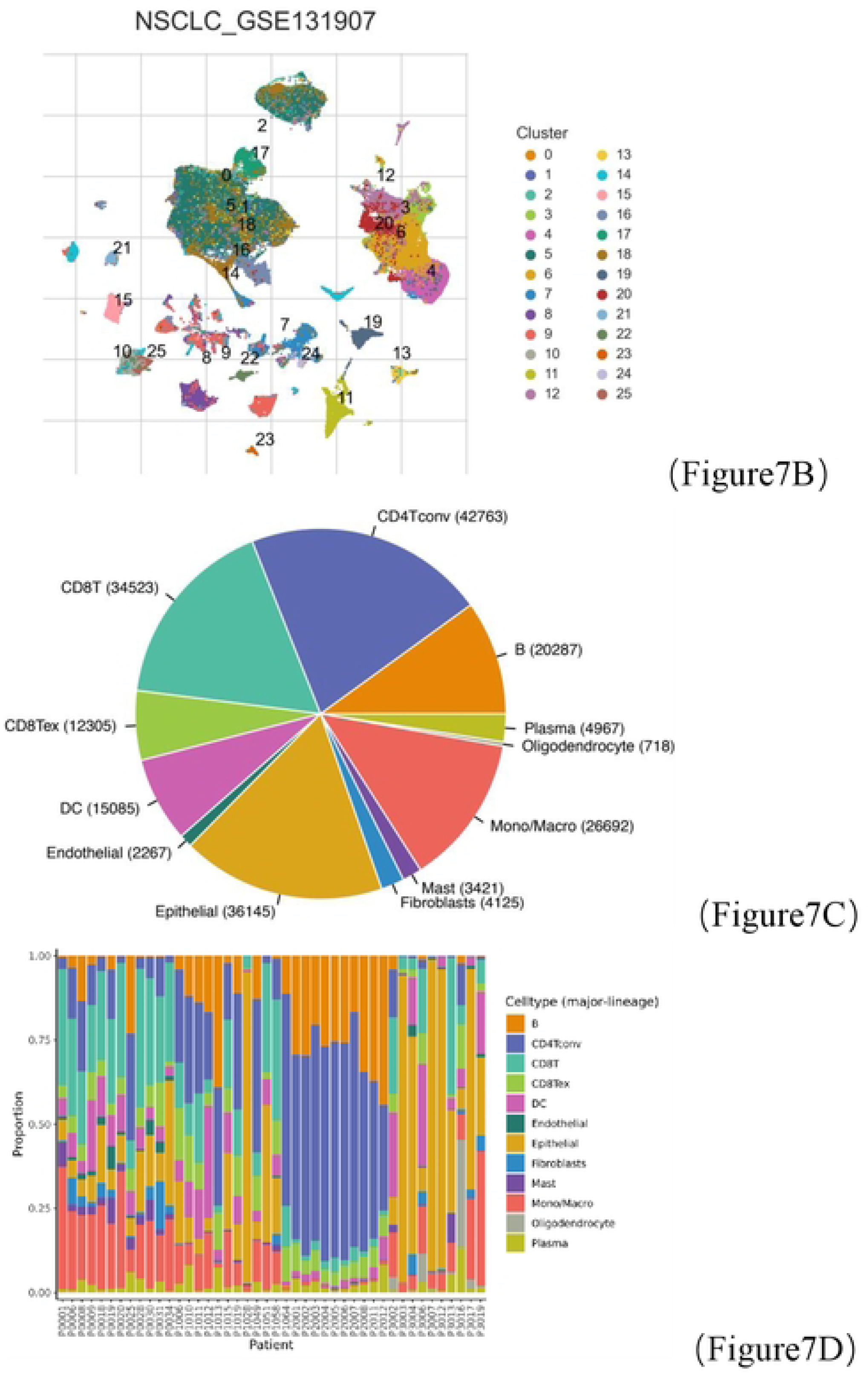
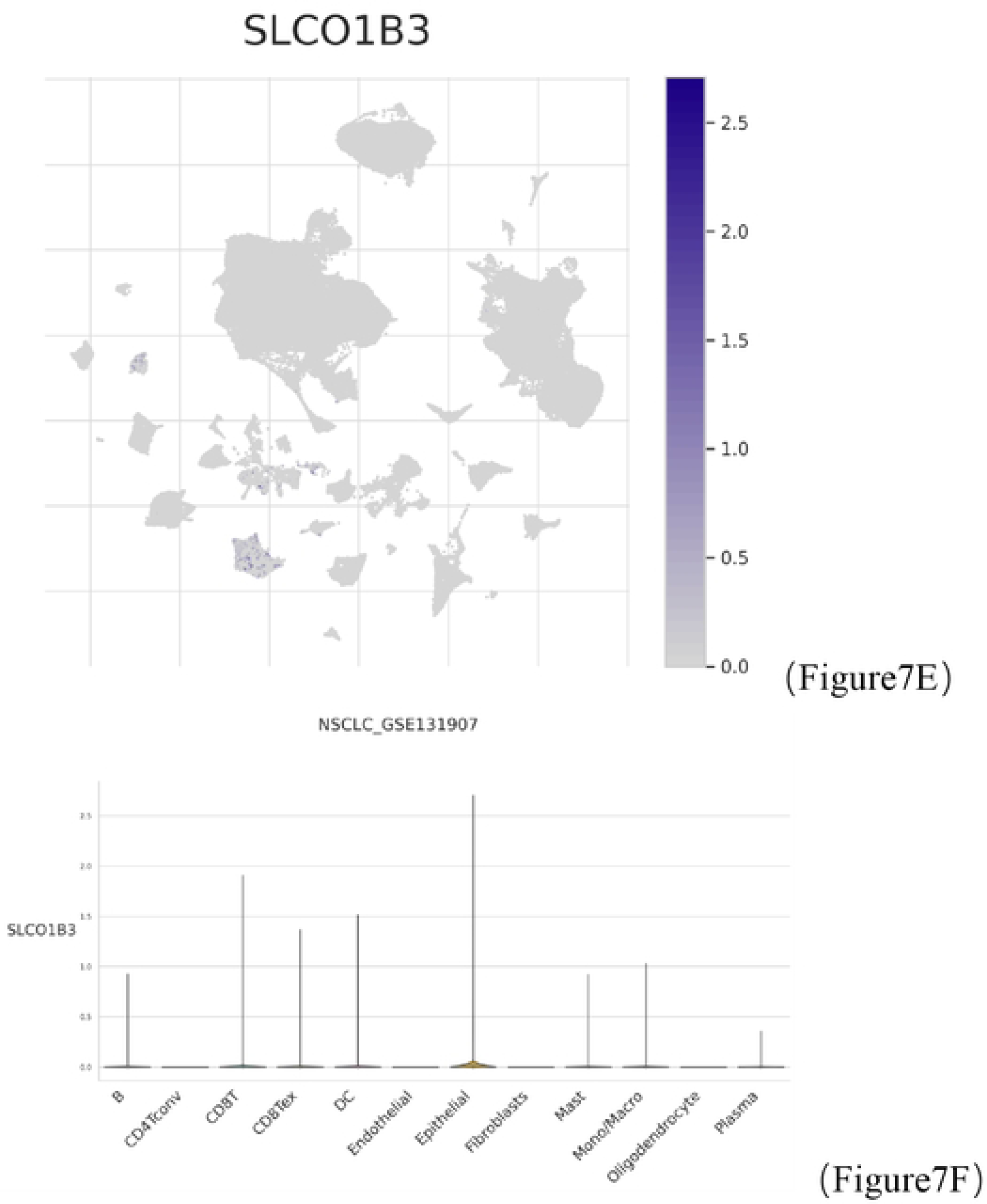
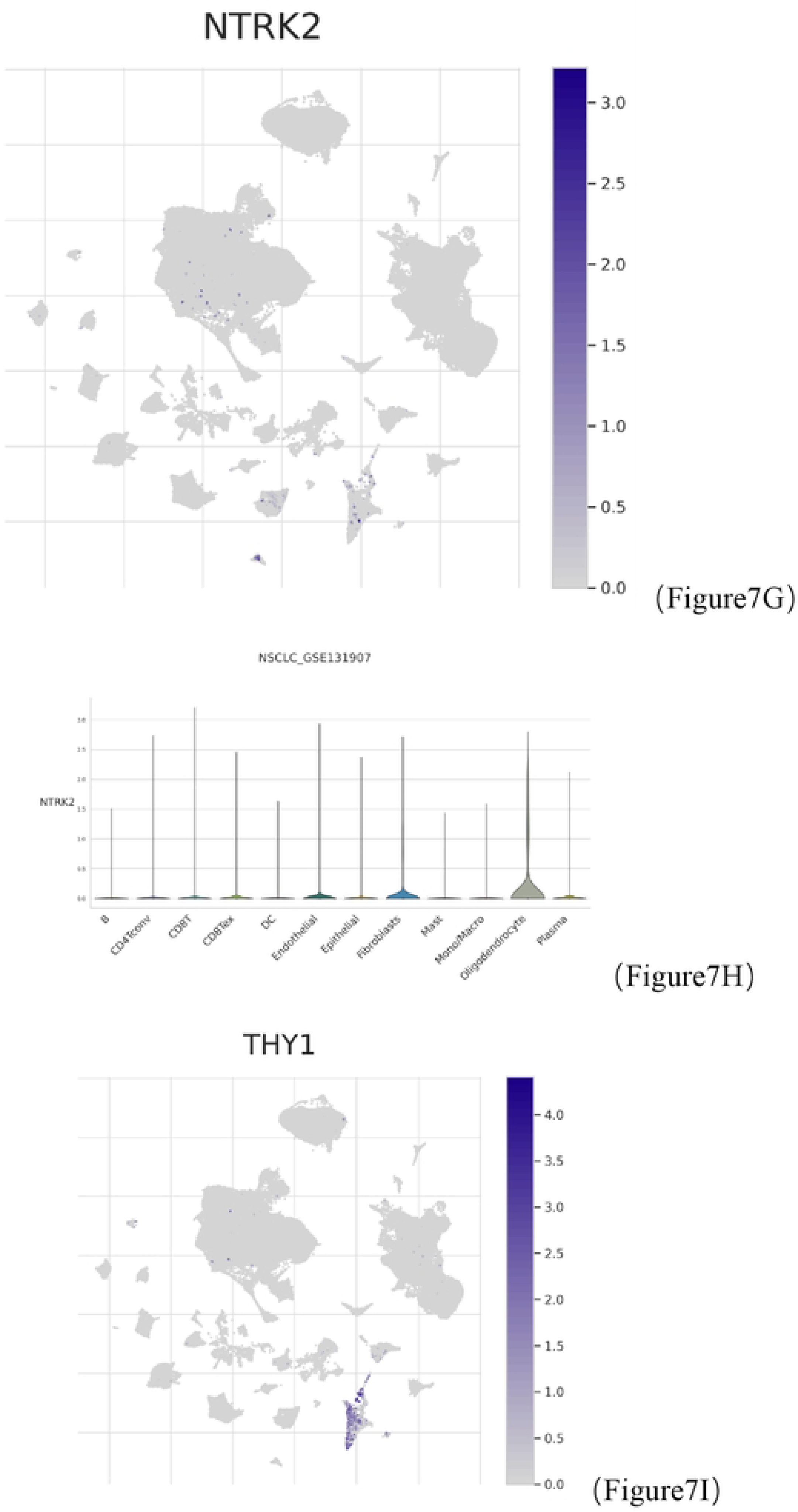
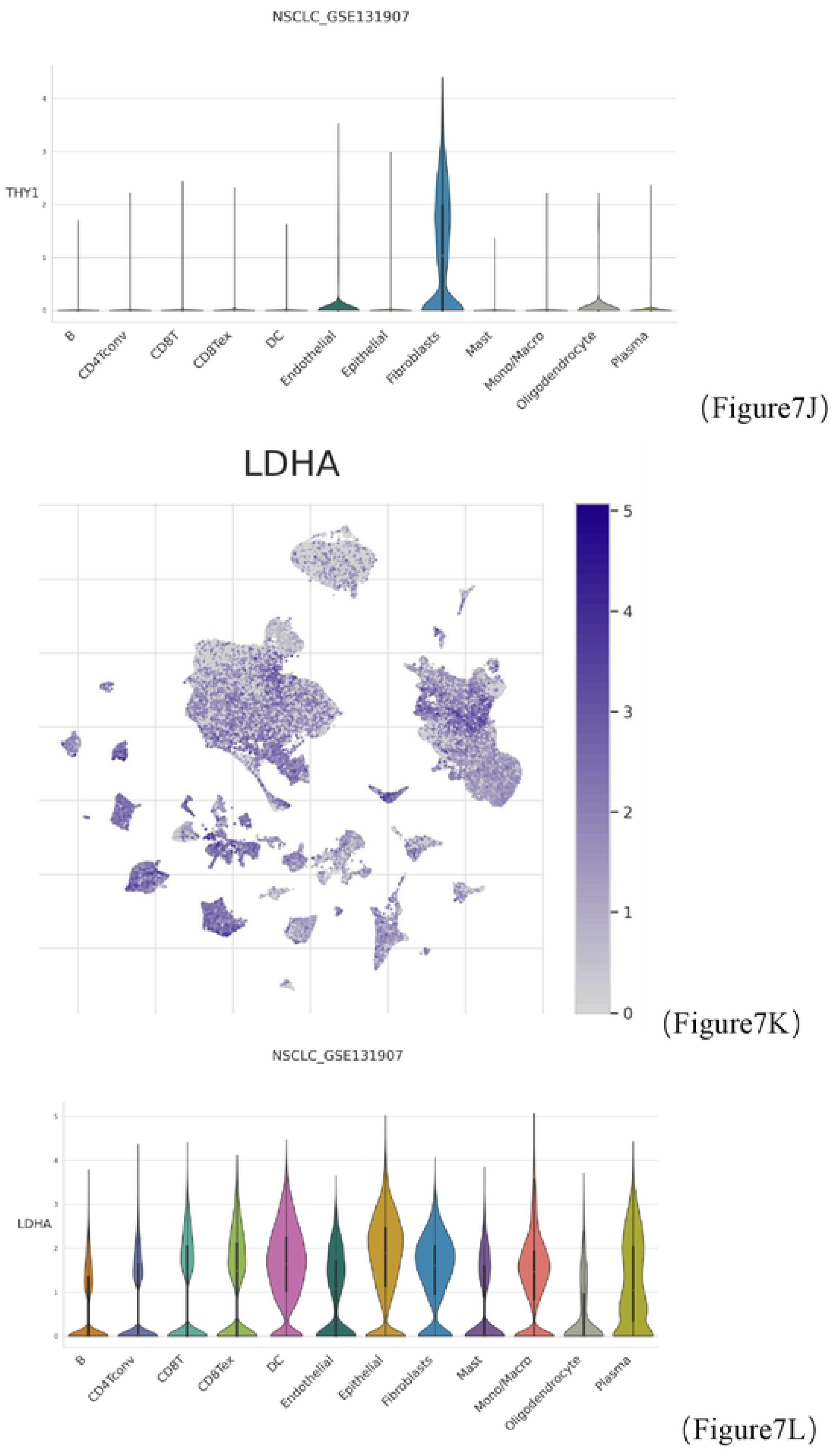
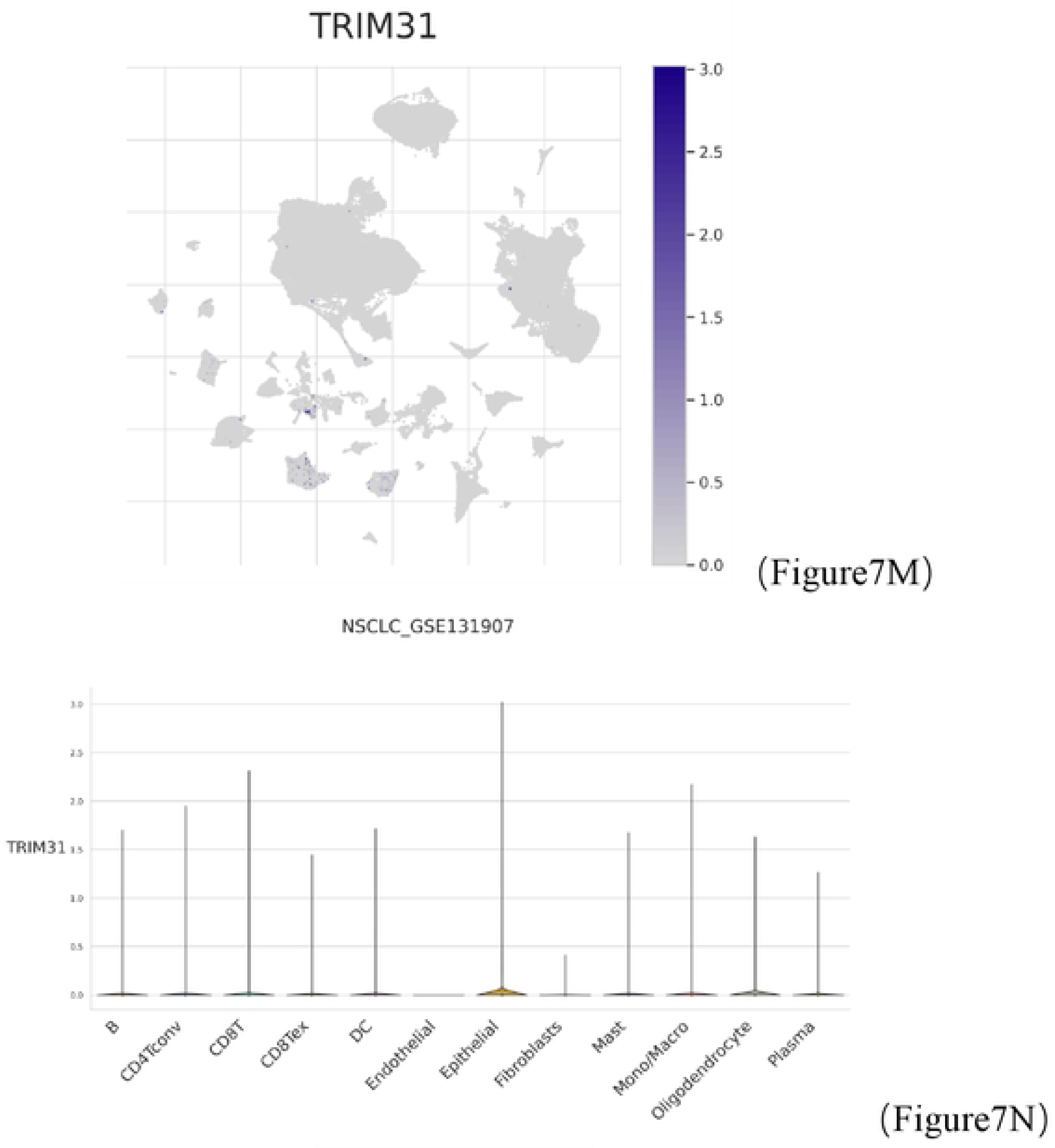
Relationship between single cell data and selected genes in non-small cell lung cancer samples. (A-B). Different clustering annotations and cell type identification. (C-D). The proportion between different cells. (E-N) Differences in the expression of selected genes between cells

### Validation of the selected genes

The expression levels of the proteins encoded by the selected genes were obtained from the HPA website. Figure 8 (A-B) shows the immunohistochemical results of the proteins encoded by LDHA. The expression results of LDHA were consistent with the results in the article, both were up-regulated in expression. The immunohistochemical results of the NTRK2-encoded protein are shown in Figure 8 (C-D), which was not detected in the tumor samples, similarly consistent with the results described in this paper. Other selected genes were not present on the network or were not significant.

**Figure 8.**
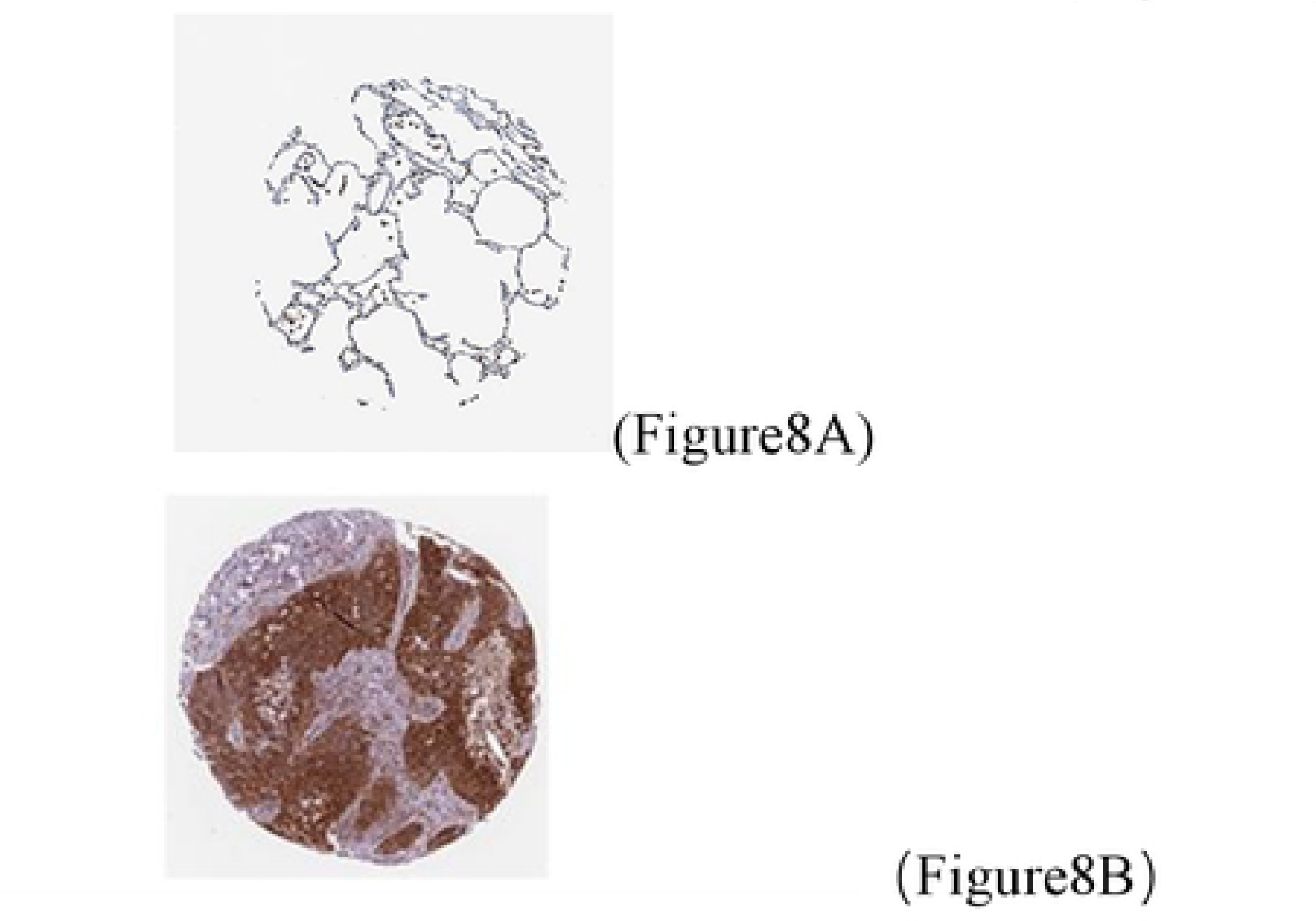
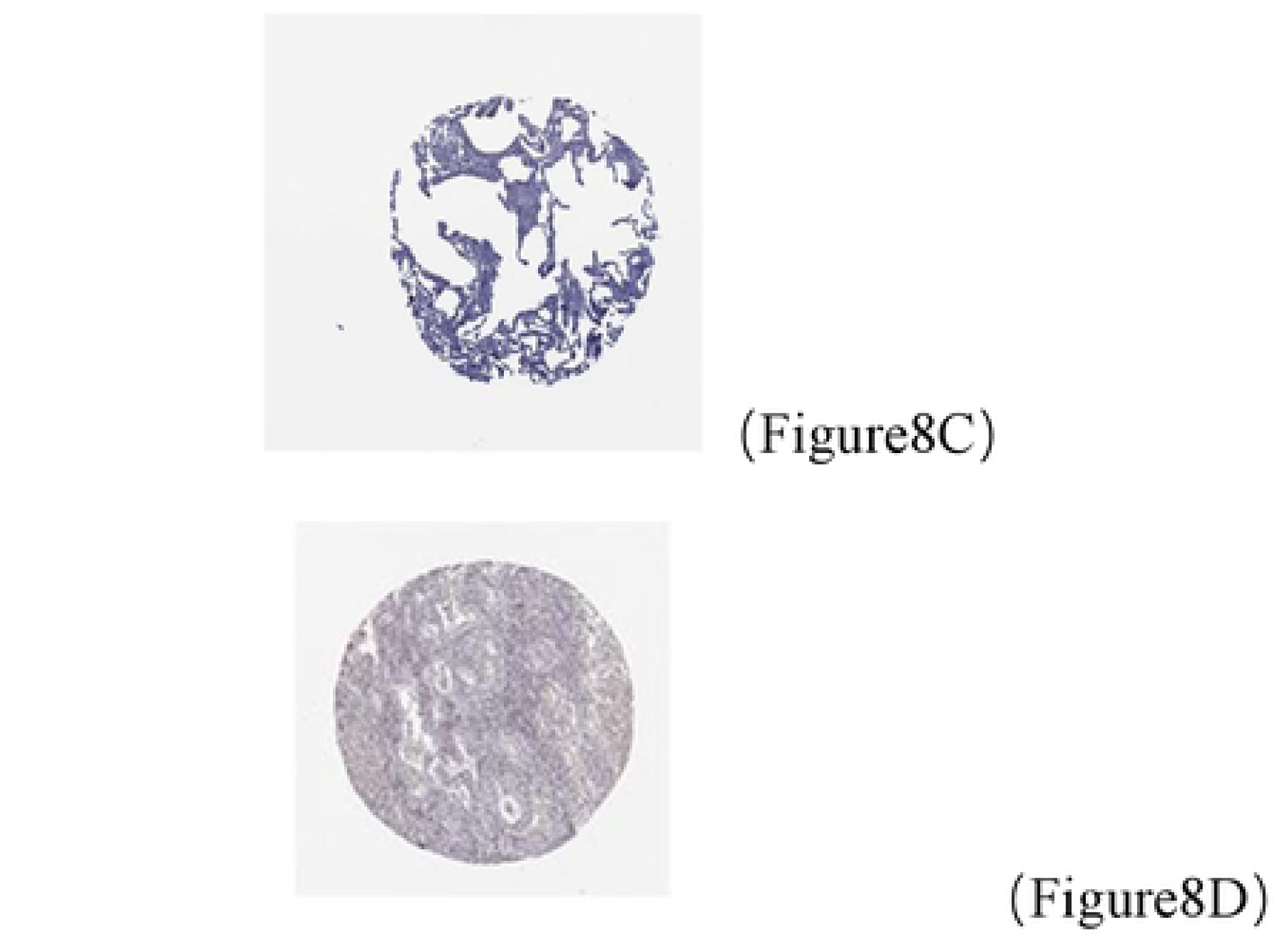
Immunohistochemical analysis. (A-B). Comparison of immunohistochemical results of the proteins encoded by LDHA between normal and tumor tissues. (C-D). Comparison of immunohistochemical results of proteins encoded by NTRK2 between normal and tumor tissues.

## DISCUSSION

Anoikis is essentially a specific mode of apoptosis that relies on interactions between cells and the ECM. Like autophagy, anoikis may play an important role in the defence of the human body. However, in the case of tumor cells, they acquire resistance to the effects of anoikis, which ultimately leads to tumor invasion, metastasis, and drug resistance [17–19].

However, there is no comprehensive and profound study on the role of anoikis genes in regulating TME, drug sensitivity and predicting patient prognosis in LA patients. After that, the relationship between selection genes and cells was compared by single cell analysis. Finally, our study was validated through the HPA website. We hope that this article will deepen the insight of apoptosis genes in lost nests and serve as a guide for future studies.

First, we obtained the anoikis genes through the website. Based on TCGA-LUAD data, a total of 129 DEGs between normal and tumor tissues were identified. After that, we further performed univariate Cox regression regression on the selected DEGs and screened out 26 genes that had an impact on patient prognosis. Then we performed consensus cluster analysis on these genes, combined with associated graphs, and finally selected the best index as 2. Also, the results of PCA analysis helped me to prove that there was a significant difference between the two groups. By plotting KM curves, there was a significant difference between ARG cluster A and B (p<0.05). After that, the DEGs between the two cluster were also compared, and most of the genes were highly expressed in cluster B. Then we compared KEGG and GO analysis between the two groups. We found that cluster B was associated with most of the tumor pathways. Also, in GO analysis, there were significant differences between the two groups. There was almost no enrichment in the A cluster in the relevant functions. Finally, GSEA analysis was performed between group A and group B. We can clearly see from the graph that group B is active in most of the pathways and functions. LASSO and multivariate Cox (multiCox) analysis for anoikis cluster-associated prognostic DEGs were conducted to establish an optimal predictive model. Five genes were finally screened: NTRK2, TRIM31, SLCO1B3, LDHA, THY1. We established the ARGs score. The train cohort was divided into high and low risk groups according to the median of the ARGs score of the training cohort. The test cohort and whole cohort were also classified into groups based on this median value. According to the test set, the training set, there is a significant difference between the KM curves of the whole data. There are significant differences in the KM curves between the test cohort, the training cohort, and the whole cohort. Also, the ROC curve based on our established ARGs score has a good performance in predicting 1, 3, and 5-year OS. All of these indicate that our ARG score has good performance in predicting prognosis.multivariate Cox regression was performed with clinical factors and risk scores together, and risk scores and advanced stage were also found to be independent of posterior factors. Heat mapping of the selected genes revealed that all were expressed in the high-risk group except NTRK2, which was expressed in the low-risk group. We plotted the alluvial diagram for all patients and we could clearly see the distribution of the sample between the different subgroups. Finally, we compared the risk scores between cluster A and B and found that there was a difference between the risk of cluster B and A, and the risk score of cluster B was higher.Combined with the patient survival curves, we found consistent findings in the text. Combined with the patient survival curves, we found consistent findings in the text. We assembled the risk scores and clinical variables together and then created Nomogram diagram. We found that Nomogram curves are also extremely valuable in predicting prognosis.The calibration curves showed that Nomogram is also highly accurate in predicting patient prognosis.Next, DCA curves were plotted for clinical factors, risk scores, and Nomogram.We found more net benefits in predicting OS at 1, 3, and 5 years.

Investigating the immune status of patients and the tumor microenvironment is our main direction. Therefore, we summarized the relationships between risk scores and immune cells. First, we analyzed the relationship between each immune subtype of cells and found a clear relationship between most of them. Then we summarized the relative content of immune cells per sample between the high- and low-risk groups. Next, we compared the differences in immune cell infiltration between the high and low risk groups and plotted violin plots. A few cells were found to differ between the two groups. We further performed an analysis of the relationship between risk scores and immune sub-cells.We also analyzed the relationship between selected genes, risk scores and immune cells, and we could see that except SLCO1B3, including risk scores and other genes and most immune cells existed in relationship. Among them, LDHA expressed in the high-risk group was closely related to DC, CD8 T cell cells, and at the same time, DC, CD8 T cells were powerful immune cells. Therefore, we believe that LDHA has an important role in regulating immune function and TME. According to existing studies, defective LDHA contributes to the immune function of CD8 T cell, which is consistent with the negative relationship results in this paper [20]. Meanwhile, the value of high LDHA expression in predicting prognosis and tumor metastasis also helps to confirm the reliability of the results in this paper [21, 22]. However, there is no report on the regulation of DC cell and TME by LDHA for the time being, which also provides a direction for future research.

Finally, we comprehensively evaluated the effect of immunotherapy and the IC50 of common drugs in patients.We can clearly see from the graph that common drugs are more sensitive in the low risk group. Meanwhile, immunotherapy was more sensitive in the low-risk group. The above results provide precise treatment direction for the current patient treatment. Then we compared the difference between the two groups of immune checkpoints, and we found a significant difference in immune checkpoints between the two groups. We downloaded single-cell data through the TISCH2 website and analyzed them, and found that LDHA was expressed significantly differently in many immune cells than the expression abundance of other genes. Also, the expression of it in DC cells was unusually abundant which helped us to verify our previous. Finally, we downloaded the immunohistochemistry results of the selected genes through the HPA website. Fortunately, we found the immunohistochemistry results of LDHA and NTRK2, and the results were consistent with this paper.

This study has the following advantages: for the first time, we analyzed the prognosis, TME and resistance of anoikis genes in LUAD patients by integrating bioinformatics analysis and single-cell data, and developed and tested predictive models.

Finally, we identified a target gene with the potential to alter the resistance of anoikis genes. This work has several limits. Data from public databases may have influenced the results. Additional clinical variables should be added to ARG_scores in order to conduct clinical studies. TME and risk score remain to be confirmed by further studies.

## Conclusion

In brief, our systematic analysis of anoikis genes revealed a comprehensive regulatory strategy that affects TME, prognosis, and clinical characteristics of LUAD patients. We also illuminate the potency of anoikis genes as biomarkers of therapeutic response. Our study reveals the clinical importance of anoikis genes and provides a valuable basis for further investigation of personalized therapy for patients with LA.

## Acknowledgements

We really appreciate the public database TCGA, GEO, TISCH2, HPA website for allowing access to their data. We declare we do not receive any third party support in conducting this research.

## Data Availability Statement

The datasets presented in this study can be found in online repositories. The names of the repository/repositories and accession number(s) can be found in the article/Supplementary Material.

## Author Contributions

Weijie Yu and Zhoulin Miao are co-first authors. All authors contributed to the concept and design of the study. Weijie Yu, Zhoulin Miao, and Bingzhang Qiao performed the data collection and analysis. Weijie Yu and Kawuli Jumai wrote the manuscript. Ilyar Sheireidin revised the manuscript. All authors commented on previous versions of the manuscript and read and approved the final manuscript.

## Ethics approval and Consent for publication

Not applicable.

